# Cell Sex and Sex Hormones Modulate Kidney Glucose and Glutamine Metabolism in Health and Diabetes

**DOI:** 10.1101/2021.08.23.457385

**Authors:** S Clotet-Freixas, O Zaslaver, C Pastrello, M Kotlyar, C McEvoy, S Farkona, A Saha, A Boshart, M Chan, M Riera, MJ Soler, A Isenbrandt, J Lamontagne-Proulx, S Pradeloux, K Coulombe, D Soulet, AB Dart, B Wicklow, JM McGavock, TD Blydt-Hansen, I Jurisica, M Woo, JW Scholey, H Röst, A Konvalinka

**Affiliations:** Toronto General Hospital Research Institute, University Health Network, Toronto, ON, Canada; Soham and Shaila Ajmera Family Transplant Centre, University Health Network, Toronto, ON, Canada; The Donnelly Centre, University of Toronto, Toronto, ON, Canada; Department of Molecular Genetics, University of Toronto, Toronto, ON, Canada; Osteoarthritis Research Program, Division of Orthopedic Surgery, Schroeder Arthritis Institute, University Health Network, Toronto, ON, Canada; Krembil Research Institute, University Health Network, Toronto, ON, Canada; Kidney Research Group, Hospital del Mar Medical Research Institute, IMIM, Barcelona, Spain; Hospital Universitari Vall d’Hebron, Division of Nephrology Autonomous University of Barcelona, Barcelona, Spain; Neurosciences Axis, CHU de Quebec Research Center - Université Laval, Québec, QC, Canada; Faculty of Pharmacy, Université Laval, Québec, QC, Canada; Department of Pediatrics and Child Health, University of Manitoba, Winnipeg, MB, Canada; Diabetes Research Envisioned and Accomplished in Manitoba Research Team, Children’s Hospital Research Institute of Manitoba, Winnipeg, MB, Canada; Department of Pediatrics, The University of British Columbia, Winnipeg, MB, Canada; Departments of Medical Biophysics and Computer Science, University of Toronto, Toronto, ON, Canada; Institute of Neuroimmunology, Slovak Academy of Sciences, Bratislava, Slovakia; Department of Medicine, Division of Endocrinology, University Health Network, University of Toronto, Toronto, ON, Canada; Department of Medicine, Division of Nephrology, University Health Network, Toronto, Canada; Department of Laboratory Medicine and Pathobiology, University of Toronto, Toronto, Canada; Institute of Medical Science, University of Toronto, Toronto, Canada

## Abstract

Male sex is a risk factor for progression of diabetic kidney disease, but the reasons for this predilection are unclear. Here, we demonstrate that cell sex and sex hormones alter the metabolic phenotype of human proximal tubular epithelial cells (PTECs). Male PTECs displayed increased glycolysis, mitochondrial respiration, oxidative stress, apoptosis, and high glucose-induced injury, compared to female PTECs. This phenotype was enhanced by dihydrotestosterone (DHT) and linked to increased mitochondrial utilization of glucose and glutamine. Studies *in vivo* pointed towards increased glutamine anaplerosis in diabetic male kidneys. Male sex was linked to increased levels of glutamate, TCA cycle, and glutathione cycle metabolites, in PTECs and in the blood metabolome of healthy youth and youth with type 2 diabetes. Conversely, female PTECs displayed increased levels of pyruvate, glutamyl-cysteine, cysteinylglycine, and a higher GSH/GSSG ratio than male PTECs, indicative of enhanced redox homeostasis. Finally, we identified transcriptional mechanisms that control kidney metabolism in a sex-specific fashion. While X-linked demethylase KDM6A facilitated metabolic homeostasis in female PTECs, transcription factor HNF4A mediated the deleterious effects of DHT in male PTECs. This work uncovers the role of sex in glucose/glutamine metabolism, that may explain the roots of sex dimorphism in the healthy and diabetic kidney.

## INTRODUCTION

Chronic kidney disease (CKD) affects >14% of the adult population and causes significant morbidity and mortality^1^. Diabetes is the leading cause of CKD worldwide, and diabetic kidney disease (DKD) is responsible for ∼40% of CKD^2^. Importantly, the cumulative incidence of DKD after 30 years of diabetes is higher in men (4.1%), compared to women (2.5%)^3^. Moreover, diabetic men are at higher risk than diabetic women of progressing to end-stage kidney disease, with hazard ratios ranging from 2.5-3.0^3–5^. Studies involving patients with type 1 or type 2 diabetes and different follow-up times (2.5-18 years) have shown that male sex is significantly associated with the development of microalbuminuria and macroalbuminuria^5–10^. However, the reasons for the sex-based nature of DKD are incompletely understood. Both clinical and animal model studies predominantly include male subjects, leaving a major gap in the understanding of the role of sex or sex hormones in kidney physiology and in DKD progression^11^. A better understanding of this sex dimorphism will lead to the development of sex-specific predictive tools and treatments for bridging this critical gap in knowledge.

Proximal tubular epithelial cells (PTECs) are the most abundant cell type in the kidney. These cells are metabolically active, as they require high amounts of energy to facilitate renal sodium reabsorption^12^. In healthy PTECs, glucose is mainly converted to pyruvate for utilization by the tricarboxylic acid (TCA) cycle, the most efficient mechanism of energy production in the form of ATP^13^. Glutamine participates in the antioxidant glutathione cycle^14^. Alterations in glucose and glutamine homeostasis are key contributors to the progression of DKD^15^. Increased glucose uptake triggers glycolysis and glutamine anaplerosis, resembling the Warburg effect^15,16^ , which is increasingly recognized in DKD^17,18^. Aberrant glucose and glutamine metabolism also activates the hexosamine pathway, leading to hypertrophy and fibrosis^19,20^. In PTECs, these metabolic alterations are linked to the development of fibrosis^21–23,24,25^. The importance of PTECs and their metabolism in DKD is supported by the remarkable effectiveness of SGLT2 inhibitors^26–30^.

We and others have demonstrated that functional and metabolic alterations in DKD develop in a sex-specific fashion^31–42^. Sex chromosomes define physiological features in adult somatic tissues^43^, and influence the development of cardiovascular disease^44^, but little is known about the effect of cell sex on kidney metabolism and disease. In diabetic mice, male sex was linked to the development of severe kidney lesions, which were attenuated in the absence of androgens^32,34^. We demonstrated that the androgen dihydrotestosterone (DHT), but not the estrogen estradiol (EST), increased enzymes involved in glucose and glutamine metabolism in PTECs. These changes were confirmed in healthy and diabetic male mice and related to kidney injury^31^. We thus postulated that energy metabolism could be the key to understand sex hormone-related effects in kidney physiology and pathophysiology.

In this study, we investigate, for the first time, the effect of cell sex on kidney metabolism. We demonstrate that male sex and androgens confer a ‘pro-oxidant’ metabolic state and accelerate high glucose-induced mitochondrial injury in PTECs. In contrast, female PTECs show decreased oxidative stress and cell death, and preserved mitochondrial function upon exposure to high glucose. We also show increased intracellular levels of glutamate, TCA, and glutathione cycle metabolites in male PTECs, while female PTECs display increased levels of pyruvate. These sex-specific differences in metabolite levels in PTECs were reproduced in adolescents with and without diabetes. At the mechanistic level, we identify hepatocyte nuclear factor 4 alpha (HNF4A), a regulatory partner of androgen receptor (AR) in the kidney^45^, as a key transcriptional regulator of glucose- and glutamine-related enzymes upregulated by androgens. We also identify X-linked lysine demethylase KDM6A as the key regulator of metabolism in the female kidney. We demonstrate that HNF4A mediates androgen-induced metabolic injury in male PTECs, while KDM6A contributes to the preservation of mitochondrial function in female PTECs. Overall, we identify a compelling link between glucose/glutamine metabolism and sex, that may explain the roots of sex dimorphism in the healthy kidney and in DKD.

## RESULTS

### Male PTECs have a more energetic and oxidative phenotype compared to female PTECs

We first investigated if sex hormones and the sex of the cell influence the metabolic phenotype of PTECs. Commercially available primary PTECs from 4 male and 5 female healthy donors of comparable ages were examined at baseline and after stimulation with sex hormones (Fig.1A).

**Figure 1.**
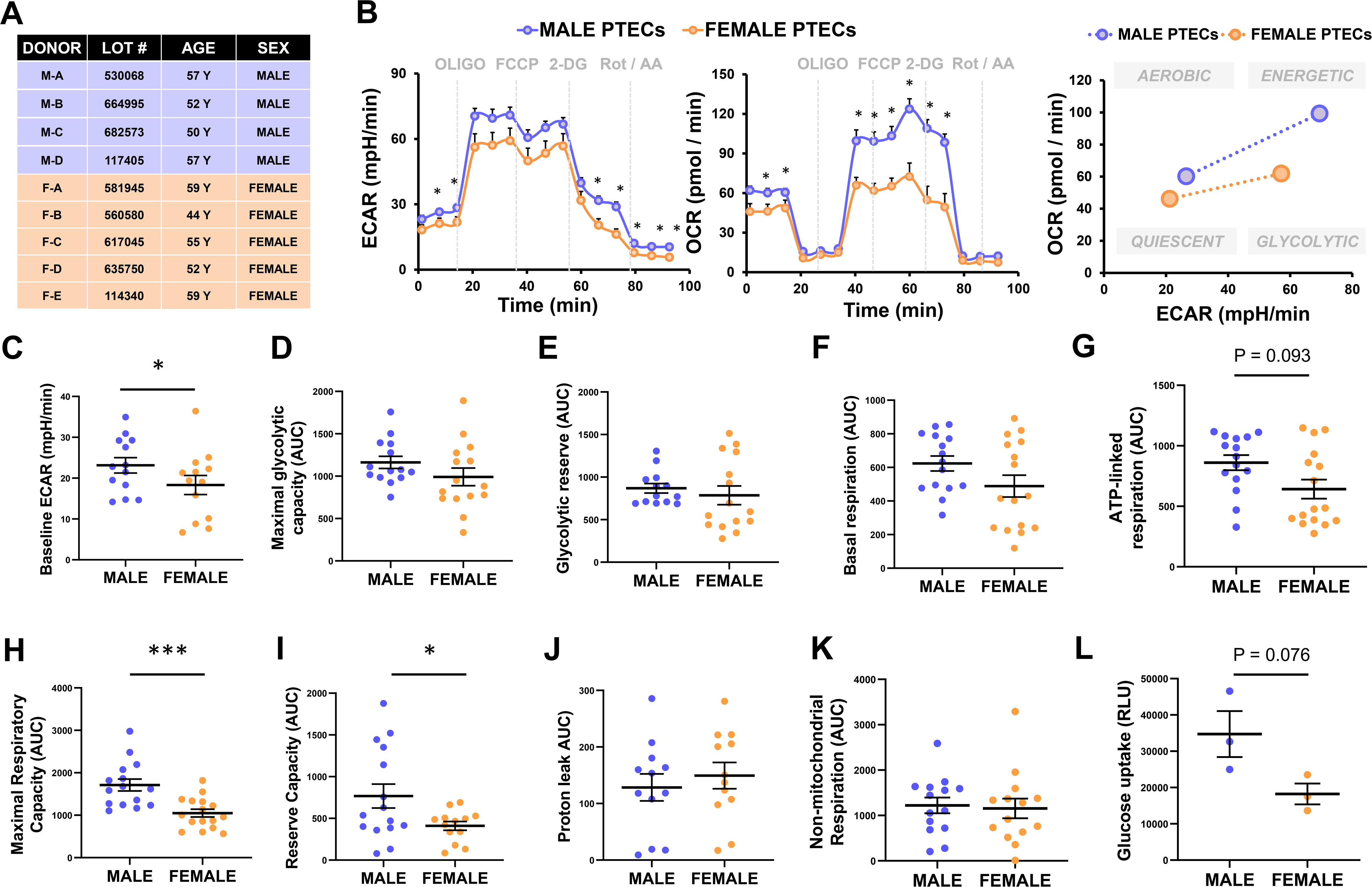
Effect of cell sex on the metabolic function of PTECs. We studied human primary PTECs from 4 male and 5 female donors (Lonza) (A). We measured glycolysis as the extracellular acidification rate (ECAR) and oxygen consumption rate (OCR) in male and female PTECs (n=3/sex; n=4-6 technical replicates/donor) in real-time in a Seahorse XFe96 analyzer. To induce metabolic stress, the following sequence of drugs was injected: 1μM oligomycin, 0.3μM FCCP, 100mM 2-DG, 1mM Rot/AA. Our results suggest a more glycolytic, oxidative and energetic phenotype in male PTECs, as compared to female PTECs (B). Baseline ECAR (C), maximal glycolytic capacity (D), and glycolytic reserve (E) were calculated from the ECAR curves in panel B. Basal respiration (F), ATP-linked respiration (G), maximal respiratory capacity (H), reserve capacity (I), proton leak (J), and non-mitochondrial respiration (K) were calculated from the OCR curves in panel B. Rapid glucose uptake (1h) was assessed in male and female PTECs by employing a luminescence assay (L) (n=3 technical replicates). *p<0.05; ***p<0.001. PTECs, proximal tubular epithelial cells; M, male; F, female; AUC, area under the curve; ECAR, extracellular acidification rate; OCR, oxygen consumption rate; OLIGO, oligomycin; FCCP, p-trifluoromethoxy carbonyl cyanide phenyl hydrazone; 2-DG, 2-deoxyglucose; Rot, rotenone; AA: antimycin A; RLU, relative luminescence units.

Male PTECs showed significantly higher glycolysis and oxygen consumption rate (OCR) than female PTECs, both at baseline and after metabolic stress (Fig.1B). We next calculated the relevant metabolic parameters from glycolysis and OCR curves (Fig.S1). Together with a significant increase in baseline glycolysis (Fig.1C), male PTECs showed a modest increment in their maximal glycolytic capacity (Fig.1D) and glycolytic reserve (Fig.1E), compared to female cells. In addition, male PTECs displayed modestly increased basal respiration (Fig.1F) and ATP-linked respiration (Fig.1G), and a significantly increased maximal respiratory capacity (Fig.1H), and reserve capacity (Fig.1I), compared to female cells. No differences between sexes were found in proton leak (Fig.1J) or non-mitochondrial respiration (Fig.1K). In conjunction with their enhanced glycolytic response, male PTECs showed a numerically higher capacity for glucose uptake, relative to female PTECs (Fig.1L).

### The metabolic effects of male sex and dihydrotestosterone are linked to increased cell injury in PTECs

We next investigated the potential role of sex hormones on the metabolic function of PTECs. In male PTECs, DHT significantly increased glycolysis and OCR at baseline, compared to CONT- and EST-treated cells. In female PTECs, both DHT and EST induced a significant increase in glycolysis and OCR at baseline and after metabolic stress, compared to CONT-treated cells (Fig.2A,B). Despite the significant effects of DHT and EST on the glycolytic activity of PTECs at baseline, none of the sex hormones significantly altered maximal glycolytic capacity or glycolytic reserve (Fig.S2A-C). In contrast, DHT significantly increased not only the basal respiration rate of male PTECs (Fig.S2D), but also their non-mitochondrial respiration (Fig.S2E). In female PTECs, the significant increase in OCR at baseline induced by DHT and EST, was accompanied by a significantly higher ATP-linked respiration (Fig.S2F), compared to CONT-treated cells.

**Figure 2.**
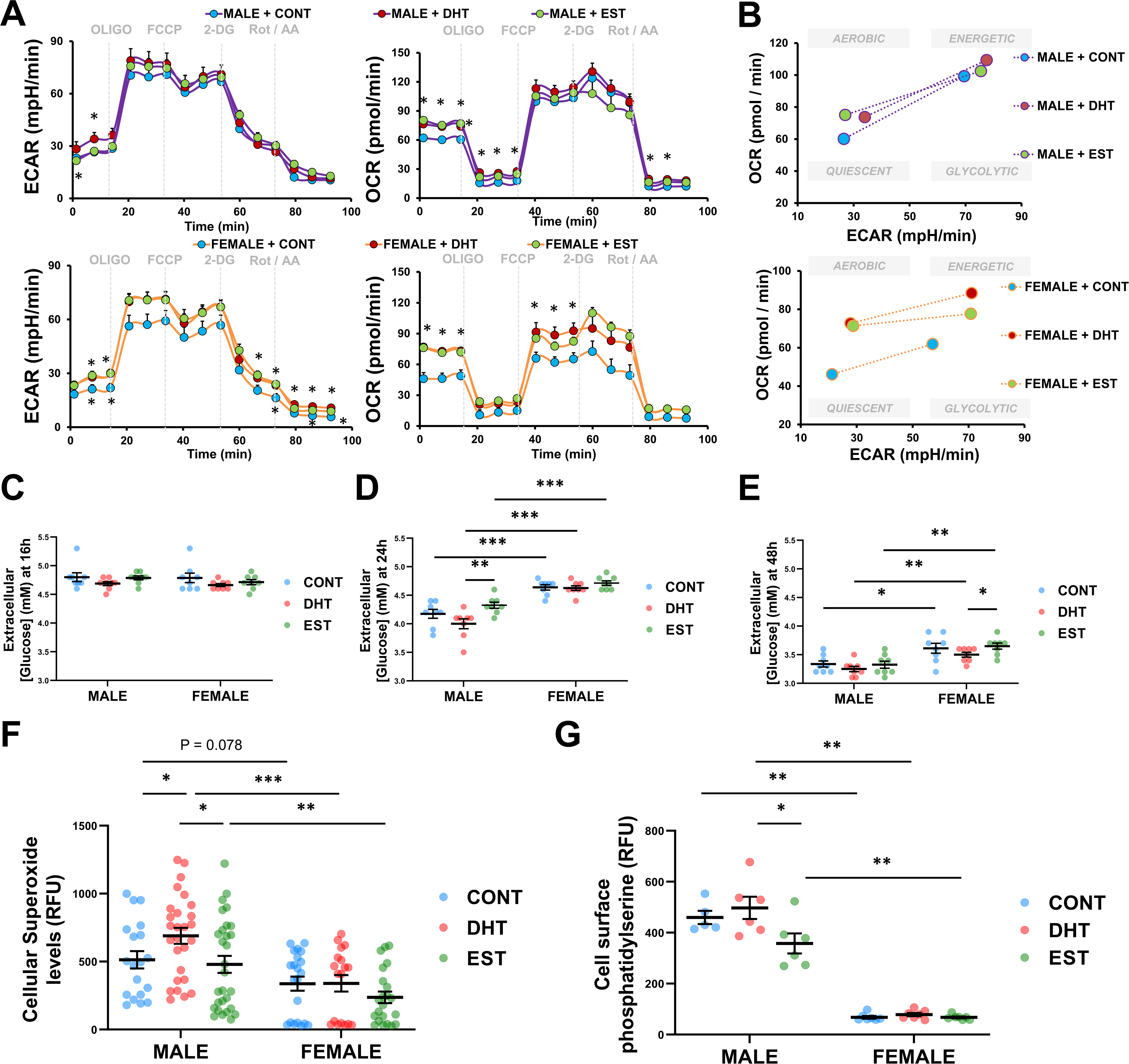
Metabolic effects of sex hormones on male and female PTECs. Glycolysis was assessed in male and female PTECs by measuring the extracellular acidification rate (ECAR) after stimulation with ethanol (CONT), 100nM DHT or 100nM EST for 16h (n=3/sex; n=4-6/treatment). Oxygen consumption rate (OCR) was also monitored (A). To induce metabolic stress, the following sequence of drugs was injected: 1μM oligomycin, 0.3μM FCCP, 100mM 2-DG, 1mM Rot/AA. The metabolic phenotype of male and female PTECs exposed to sex hormones was evaluated by plotting ECAR in the X axis, and OCR in the Y axis (B). Glucose levels in the media were assessed using Accu-check Aviva Nano strips in CONT-, DHT-, and EST-treated male and female PTECs at 16h (C), 24h (D), and 48h (E) (n=2/sex; n=4-6/treatment). After 16h of hormone exposure, intracellular levels of superoxide ion were measured to assess oxidative stress(F) (n=3/sex; n=4-6/treatment), and surface levels of phosphatidylserine were measured to assess early apoptosis (G) (n=2/sex; n=4-6/treatment). *p<0.05; **p<0.01; ***p<0.001. PTECs, proximal tubular epithelial cells; CONT, control; DHT, dihydrotestosterone; EST, 17β-estradiol; ECAR, extracellular acidification rate; OCR, oxygen consumption rate; FCCP, p-trifluoromethoxy carbonyl cyanide phenyl hydrazone; 2-DG, 2-deoxyglucose; Rot, rotenone; AA: antimycin A; RFU, relative fluorescence units; PS, phosphatidylserine.

We next examined if the functional differences linked to cell sex and sex hormones in PTECs related to a different propensity to consume glucose. We did not observe any significant differences in glucose levels in the media, at 16h (Fig.2C). However, at 24h and 48h, all groups of male PTECs displayed a significant reduction in the extracellular levels of glucose (indicating increased consumption), compared to female PTECs receiving the same treatment (Fig.2D-E). Moreover, DHT significantly enhanced the glucose consumption in male and female PTECs, compared to CONT and EST treatments (Fig.2D,E).

In metabolic disorders such as DKD, excessive glucose consumption and utilization in the TCA cycle may result in mitochondrial stress and cell death^46^. We thus evaluated if the observed changes in metabolic function and glucose consumption induced by cell sex and sex hormones in PTECs were related to altered oxidative stress and apoptosis^15,20,47^. Importantly, increased glucose and oxygen consumption in male PTECs were accompanied by significantly higher levels of oxidative stress (Fig.2F) and early apoptosis (Fig.2G), compared to female cells. DHT-treated male PTECs showed significantly higher oxidative stress levels than CONT-treated cells (Fig.2F), as well as significantly augmented early apoptosis compared to EST-treated cells (Fig.2G). Despite increasing OCR in male PTECs and altering the metabolic function of female PTECs (Fig.2A,B), EST treatment did not increase oxidative stress or apoptosis, regardless of the cell sex (Fig.2F,G).

### Androgen receptor mediates metabolic perturbations in male PTECs

DHT had a dominant effect on glucose consumption in both male and female PTECs, and promoted glycolysis, oxidative stress, and apoptosis in male PTECs (Fig.2). To elucidate if the metabolic actions of DHT were mediated by AR, we exposed male and female PTECs to anti-androgen treatment for 2h prior to sex hormone stimulation. Two different AR inhibitors with distinct mechanisms of action were employed: flutamide^48^ and enzalutamide^49^ (Fig.S3A). Pre-treatment for 2h with either FLUT (1µM) or ENZ (1µM) significantly prevented the DHT-induced increase in glycolysis (Fig.S3B, D-F, I) and mitochondrial respiration (Fig.S3C, G-I) in male PTECs and, to a lesser extent, in female PTECs. Pre-treatment with FLUT also numerically reduced superoxide ion levels in DHT-treated male and female PTECs. A similar trend was observed in DHT-treated male and female PTECs pre-treated with ENZ (Fig.S3J). Overall, the effects of AR inhibition were greater in male PTECs than in female PTECs. This sex dimorphism could be explained by increased AR expression in male PTECs (Fig.S3K). These data suggest that DHT-induced metabolic alterations in PTECs are mediated by AR.

### High glucose-induced mitochondrial dysfunction and ATP depletion are accelerated in male PTECs

We next investigated if high glucose is a modifier of the identified sex differences in the metabolism of PTECs. Male and female PTECs were exposed to high glucose for 2h (mimicking acute hyperglycemia), and 48h-96h (chronic hyperglycemia).

In male PTECs, high glucose exposure resulted in a marked increase in glycolysis and mitochondrial respiration at 2h (Fig.3A). While the gradual increase in mitochondrial respiration persisted after 48h of hyperglycemia, the basal glycolytic rate of male PTECs started to decline at this time point. After 96h of exposure to high glucose, male PTECs displayed a reduction in both glycolysis and OCR, as compared to the previous time point. In female PTECs, 2h of hyperglycemia was also associated with a significant increase in glycolysis and OCR. Interestingly, both parameters were further increased at 48h, especially after metabolic stress. At 96h, high glucose induced a more pronounced increase in OCR in female PTECs, but a reduction in the glycolytic rate, as compared to the previous time points (Fig.3A).

**Figure 3.**
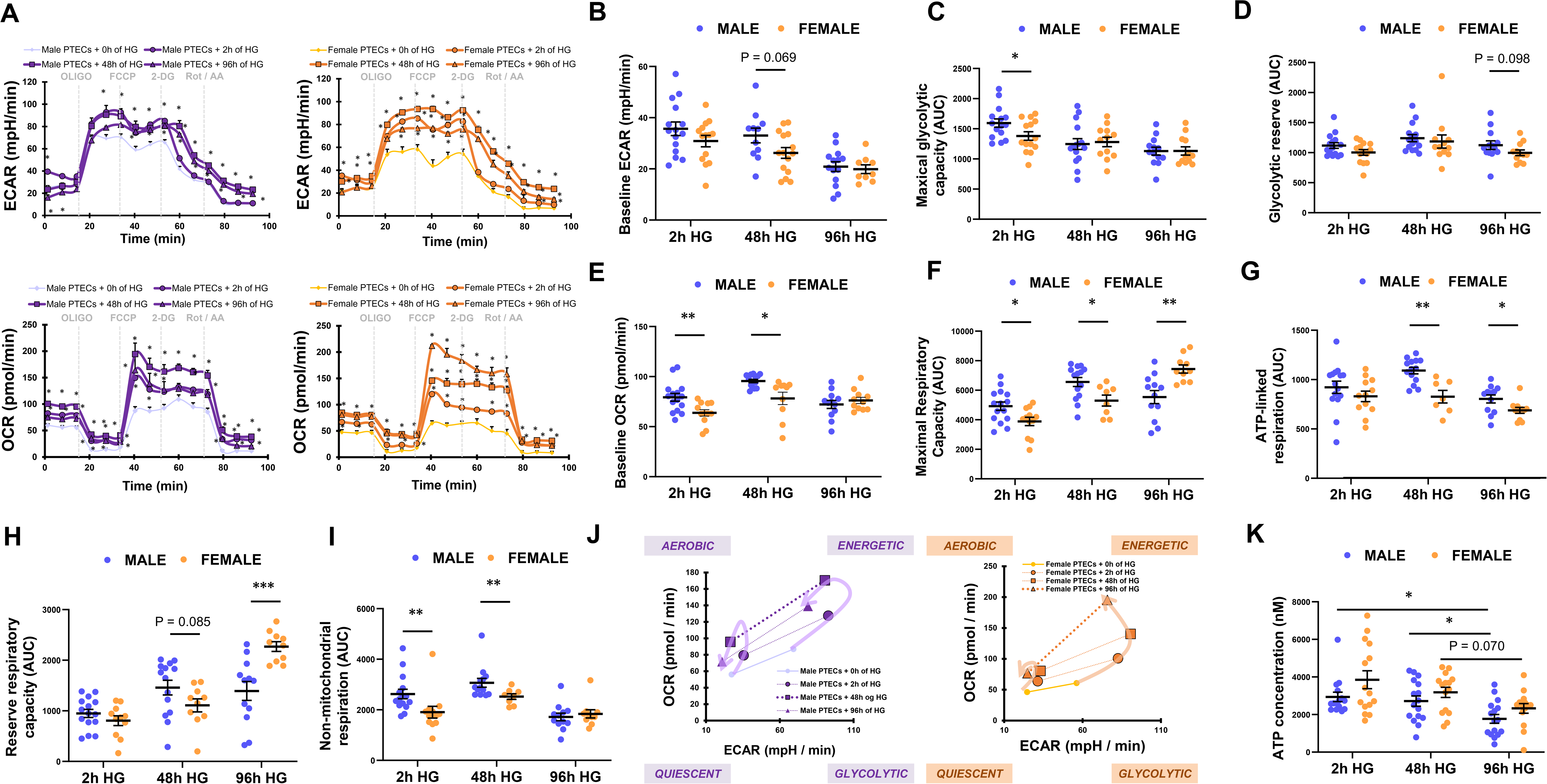
Effects of acute and chronic hyperglycemia on the metabolism of male and female PTECs. ECAR and OCR were monitored to assess glycolysis and mitochondrial function, respectively, in male and female PTECs at baseline and after 2h, 48h, and 96h of exposure to hyperglycemia (25mM glucose); (n=2/sex; n=4-6/time point) (A). To induce metabolic stress, the following sequence of drugs was injected: 1μM oligomycin, 0.3μM FCCP, 100mM 2-DG, 1mM Rot/AA. Baseline ECAR (B), maximal glycolytic capacity (C), and glycolytic reserve (D) were calculated from the ECAR curves in panel A. Baseline OCR (E), maximal respiratory capacity (F), ATP-linked respiration (G), reserve respiratory capacity (H), and non-mitochondrial respiration (I) were calculated from the OCR curves in panel A. The evolution of the metabolic phenotype of male and female PTECs during hyperglycemia was visualized by plotting ECAR on the X axis and OCR on the Y axis (J). The intracellular levels of ATP were also measured (K). *p<0.05; **p<0.01; ***p<0.001. PTECs, proximal tubular epithelial cells; AUC, area under the curve; ECAR, extracellular acidification rate; OCR, oxygen consumption rate; FCCP, p-trifluoromethoxy carbonyl cyanide phenyl hydrazone; 2-DG, 2-deoxyglucose; Rot, rotenone; AA: antimycin A.

The differences between male and female PTECs exposed to high glucose over time were reflected in glycolysis- and OCR-related metabolic parameters. After 2h and 48h of high glucose, male PTECs still showed a higher baseline glycolysis (Fig.3B), maximal glycolytic capacity (Fig.3C), glycolytic reserve (Fig.3D), basal respiration (Fig.3E), maximal respiratory capacity (Fig.3F), ATP-linked respiration (Fig.3G), reserve respiratory capacity (Fig.3H) and non-mitochondrial respiration (Fig.3I) than female PTECs. This effect was not observed after 96h of high glucose exposure. At this time point, the maximal respiratory capacity (Fig.3F) and reserve respiratory capacity (Fig.3H) were significantly higher in female, compared to male PTECs.

Our findings suggest that high glucose conditions elicit sex-specific metabolic changes in PTECs, that are time-dependent. Male PTECs displayed a more energetic phenotype during acute exposure (2h), and a switch toward a more quiescent state at 48h, with a clear reduction in their glycolytic rate and mitochondrial activity at 96h. This slowing of glycolytic and oxygen consumption rates was delayed in female PTECs, which showed a more energetic phenotype after 48h of high glucose, and a change towards a more oxidative phenotype at 96h (Fig.3J). High glucose induced a decline in the intracellular levels of ATP over time. Again, this effect was more pronounced in male PTECs, which showed the lowest ATP levels at 96h (Fig.3K). These data suggest that male sex accelerates high glucose-induced reduction in glycolysis and OCR, and subsequent ATP depletion in PTECs.

We also investigated if co-treatment with sex hormones modified the effects of high glucose on PTEC metabolic function (Fig.S4). DHT-treated PTECs displayed the highest levels of ATP and oxidative stress after 96h of high glucose, and this effect was more pronounced in male compared to female PTECs. In line with their reduced respiration, EST-treated male PTECs exposed to high glucose showed lower levels of ATP and superoxide ion, compared to CONT- and DHT-treated cells (Fig.S4J,K). These data suggest that EST accelerates the decline in mitochondrial function induced by high glucose in male PTECs, while delaying this effect in female PTECs. In contrast, DHT promotes oxidative stress in the setting of high glucose, particularly in male PTECs.

### Increased mitochondrial utilization of glucose and glutamine contributes to the enhanced metabolic activity of male PTECs

Early alterations in the metabolism of glucose, glutamine and pyruvate are key contributors to DKD^15,20^. We therefore deciphered the specific contribution of these substrates to the metabolic phenotype of PTECs. We subjected male and female PTECs to nutrient restricted conditions, prior to the measurement of glycolysis and OCR. As observed under non-restricting conditions (Fig.1), exposure to glucose alone resulted in a similar extent of increased respiration and glycolysis in male compared to female cells (Fig.4A-C). Importantly, when glutamine was the only substrate present in the media, male PTECs showed the greatest increase in their maximal respiratory capacity and respiratory reserve, compared to female PTECs (Fig.4D-F). In contrast, no marked sex differences in OCR and glycolysis were observed in the presence of pyruvate (Fig.4G-I).

**Figure 4.**
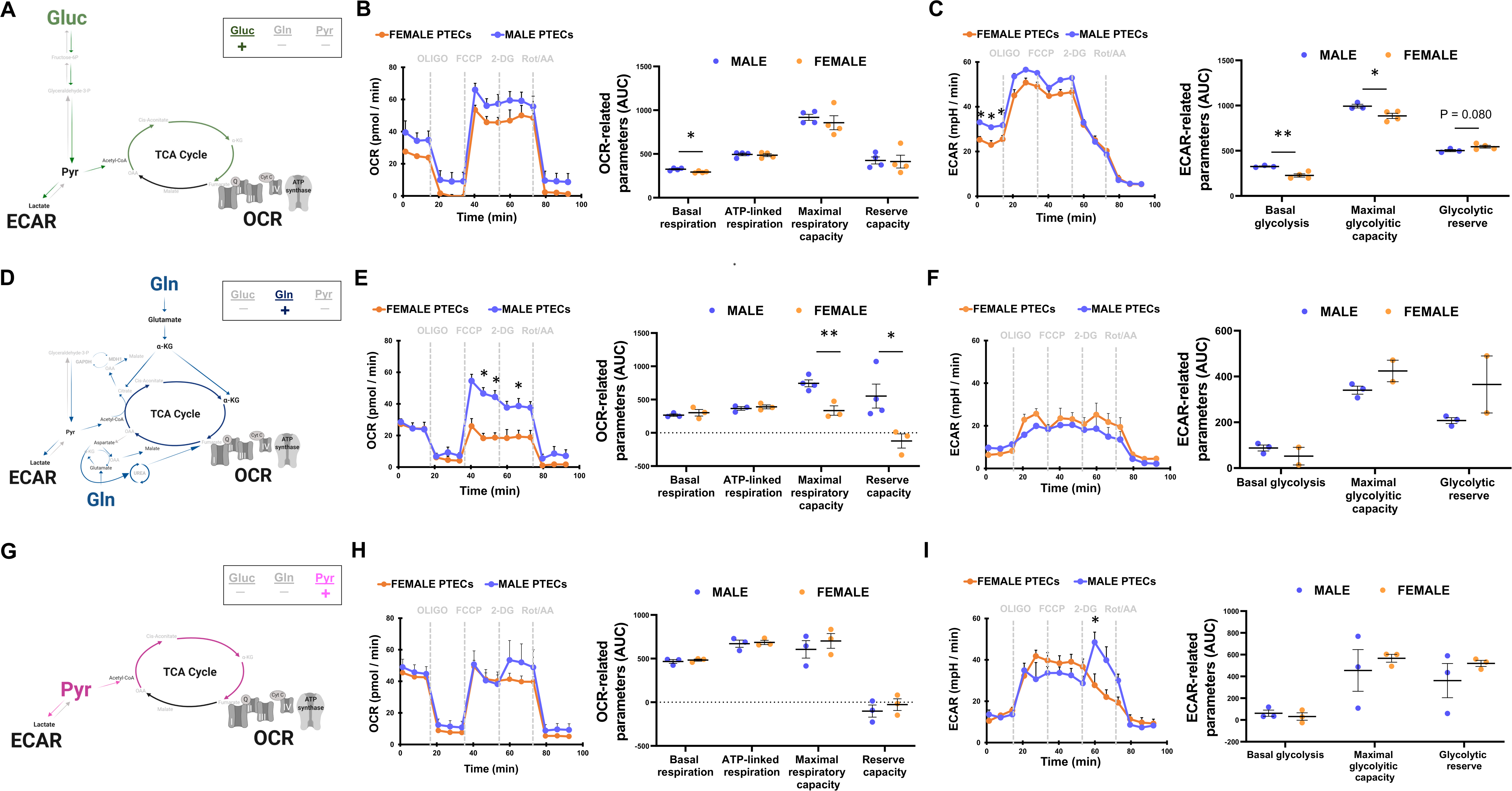
Effects of glucose, glutamine, and pyruvate on the sex-based differences in the metabolic function of PTECs. Oxygen consumption rate (OCR) and glycolysis (ECAR) were measured in male and female PTECs exposed to glucose-only (A-C), glutamine-only (D-F) or pyruvate-only (G-I) conditions for 1h (n=2-4). To induce metabolic stress, the following sequence of drugs was injected: 1μM oligomycin, 0.3μM FCCP, 100mM 2-DG, 1μM Rot/AA. Basal respiration, ATP-linked respiration, maximal respiratory capacity, and reserve respiratory capacity were calculated from the OCR curves. Basal glycolysis, maximal glycolytic capacity, and glycolytic reserve were calculated from the ECAR curves. *p<0.05; **p<0.01; ***p<0.001. PTECs, proximal tubular epithelial cells; Gluc, glucose; Gln, glutamine; Pyr, pyruvate; α-KG, alpha-ketoglutarate; TCA, tricarboxylic acid; AUC, area under the curve; ECAR, extracellular acidification rate; OCR, oxygen consumption rate; FCCP, p-trifluoromethoxy carbonyl cyanide phenyl hydrazone; 2-DG, 2-deoxyglucose; Rot, rotenone; AA: antimycin A. Illustrations in panels A, D, and G were created with BioRender.com.

Our substrate-restricting studies suggest a higher mitochondrial utilization of glucose and glutamine by male PTECs, compared to female PTECs. In turn, female PTECs showed a relative preference for pyruvate utilization over glucose and glutamine. In male PTECs, glucose seems to be more rapidly metabolized through glycolysis, and consequently, glutamine may be supporting the mitochondrial TCA cycle to a higher extent than in female PTECs.

### Male sex is linked to increased circulating levels of glucose, lactate, glutamine, and glutamate in diabetic and healthy mice

Metabolites are considered the final output of the cell biological activity and reflect the metabolic states of a living system^50^. Whereas lactate is the end-product of glucose utilization through glycolysis, glutamate is the product of glutaminolysis and a key intermediate for the entry of glutamine-derived carbons to the TCA cycle in the mitochondria, after further conversion to alpha-ketoglutarate (α-KG) (Fig.5A)^51,52^. We next examined *in vivo* the circulating and urinary levels of glucose, glutamine, and their breakdown products (lactate and glutamate).

**Figure 5.**
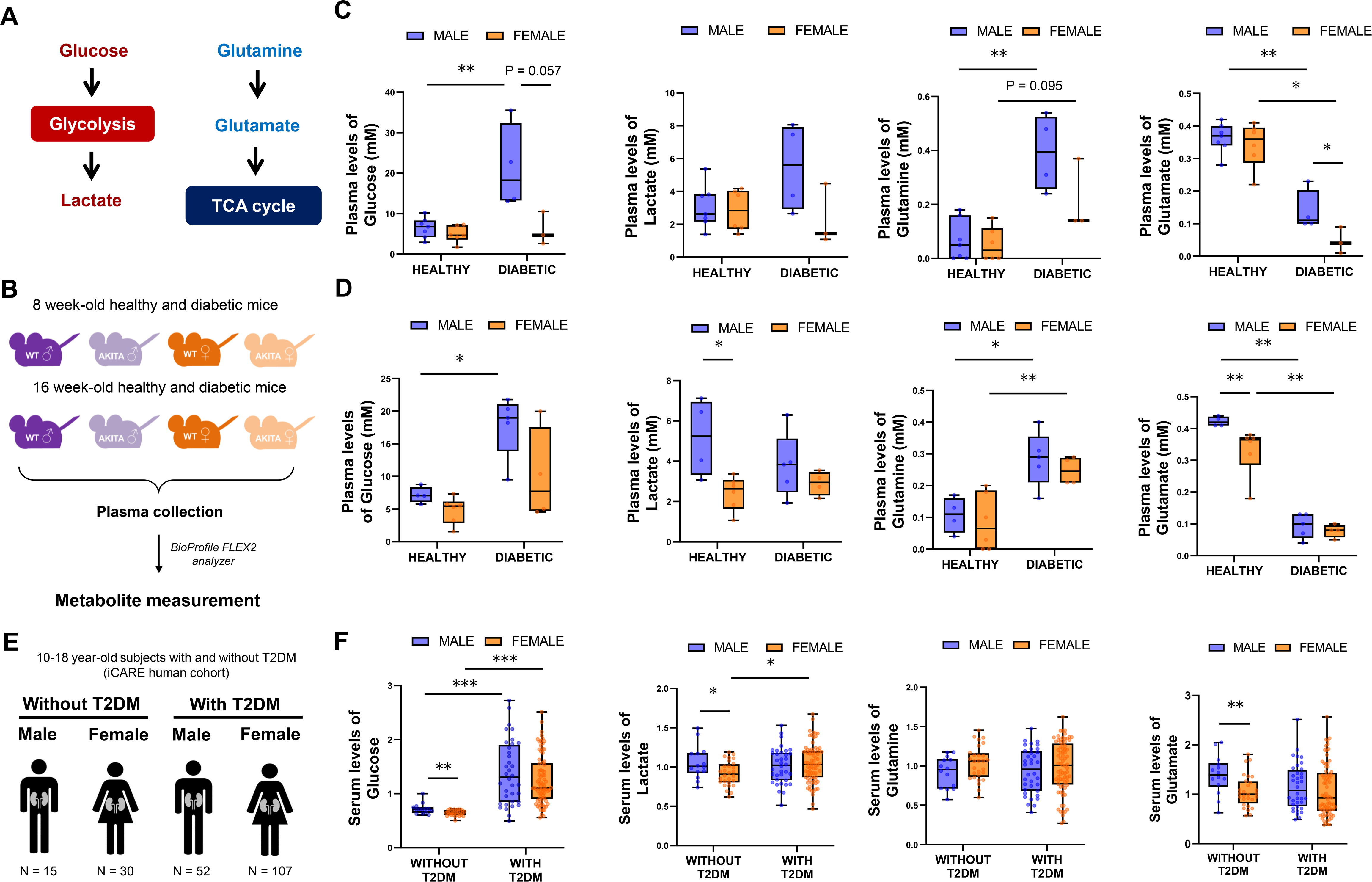
Sex differences in glucose and glutamine metabolism *in vivo* and in humans. Glucose can generate lactate via glycolysis, while glutamine can be converted to glutamate, which can enter the TCA cycle (A). Glucose, lactate, glutamine and glutamate were measured using a BioProfile® FLEX2™ Analyzer in the plasma of male and female type 1 diabetic (Akita) mice, and their healthy wild-type littermates (B). Two different age groups were studied: 8 weeks (C), and 16 weeks (D) (n=3-7 animals/group). Sex differences in glucose, lactate, glutamine, and glutamate levels were also examined in the serum of men without T2DM (n=15), women without T2DM (n=27), men with T2DM (n=47), and women with T2DM (n=91) (E-F). Data normalization was performed to correct variation emerging from inter-day tuning differences on instrument performance. Each compound was corrected in run-day blocks by registering the medians to 1.00 and normalizing each data point proportionately. Metabolite measurements are thus presented as normalized mass spectrometry-based intensities. *p<0.05; **p<0.01; ***p<0.001. PTECs, proximal tubular epithelial cells; CONT, control; DHT, dihydrotestosterone; EST, 17β-estradiol; WT, wild-type; α-KG, alpha-ketoglutarate; TCA, tricarboxylic acid; T2DM, type 2 diabetes mellitus.

We first measured these 4 metabolites in the plasma of control and type 1 diabetic (T1DM) male and female Akita mice. At 16 weeks of age, Akita mice show microalbuminuria and glomerular enlargement, indicative of early DKD^53^. We examined sexually mature animals at two different ages: 8 weeks and 16 weeks^54^ (Fig.5B). Hyperglycemia in diabetic males was associated with increased lactate levels at 8 weeks, but not at 16 weeks of age. Diabetic females showed no changes in plasma lactate levels, compared to healthy animals. All diabetic groups showed a clear increase in the plasma levels of glutamine and a significant reduction in glutamate, compared to their healthy controls, and regardless of sex and age (Fig.5C,D).

Male sex was associated with increased circulating levels of glucose, lactate, glutamine, and glutamate. This effect was modified by age and diabetes. The circulating levels of lactate and glutamate were significantly higher in 16-week-old healthy males, compared to females. In contrast, within the diabetic groups, sex differences in the plasma levels of glucose, lactate, glutamine, and glutamate were more evident at 8 weeks than at 16 weeks (Fig.5C,D). Interestingly, male mice showed lower urinary excretion of glutamine and glutamate than female mice, and this decrease was prevented by androgen reduction through gonadectomy (Fig.S5A,B). These data suggest that male sex and androgens promote the utilization of glutamate by kidney cells, rather than its excretion.

### The link between male sex and increased circulating levels of glucose, lactate, and glutamate is conserved in humans

To test the translational potential of our findings, we investigated the effects of sex and diabetes on the serum metabolome of 10-18 years old adolescents (iCARE cohort)^55^ with early onset type 2 diabetes (T2DM, n=159), and their age- and body weight-matched controls without diabetes (n=45) (Fig.5E). This cohort was particularly relevant, as our findings *in vivo* supported sex-specific effects early in diabetes. The median duration of diabetes was 2.3 years and was comparable between male and female patients (Table 1). On average, patients with T2DM displayed a significant but minimal increase in urine albumin-to-creatinine ratio (median 0.7 mg/mmol), compared to controls without T2DM (median 0.2 mg/mmol). Hypertension was present in 50% of patients with T2DM and 40% of controls. Patients with T2DM showed an increase in eGFR (143.1 ml/min/1.73m^2^ vs 139.4 ml/min/1.73m^2^), indicative of early hyperfiltration. This eGFR increase was more marked in males with T2DM (median 154.8 ml/min/1.73m^2^) compared to females (median 139.4 ml/min/1.73m^2^). Overall, these clinical features are indicative of early stages of DKD among both male and female patients with T2DM.

**Table 1.** Demographics of adolescents with type 2 diabetes and matched controls. Group-to-group differences in age at baseline, duration of diabetes, HbA1c, BMI, ACR, and eGFR were calculated using the Mann-Whitney test, and data are expressed as median (interquartile range). The eGFR was calculated using the iCARE equation (Dart, *et al., Pediatr Nephrol*. 2019): 50.7 × BSA^0.816^ × (height (cm)/creatinine)^0.405^ × 0.8994 if sex = female | 1 otherwise. Group-to-group differences in the frequency of participants with hypertension were calculated using the Fisher exact test. Abbreviations: T2DM, type 2 diabetes mellitus; HbA1c, glycated hemoglobin; BMI, body mass index; ACR, albumin to creatinine ratio; eGFR, estimated glomerular filtration rate; N/A, not available.

|  | Controls without T2DM | Patients with T2DM | P value |
| --- | --- | --- | --- |
| **N** | 45 | 159 | N/A |
| **Sex (% Female)** | 66.67 | 67.30 | 1.000 |
| **Age at Baseline (years)** |  |  |  |
| Total | 16.87 (13.77, 18.10) | 14.95 (13.14, 16.65) | 0.020 |
| Male | 14.99 (13.60, 18.26) | 15.48 (13.43, 16.75) | 0.383 |
| Female | 16.91 (14.28, 17.86) | 14.67 (13.09, 16.51) | 0.029 |
| **Duration of diabetes (years)** |  |  |  |
| Total | N/A | 2.28 (1.16, 4.05) | N/A |
| Male | N/A | 2.57 (1.27, 3.92) | N/A |
| Female | N/A | 1.96 (1.05, 4.05) | N/A |
| **HbA1c (%)** |  |  |  |
| Total | 5.60 (5.50, 5.80) | 9.50 (7.05, 11.45) | <0.001 |
| Male | 5.50 (5.50, 5.70) | 9.45 (6.87, 11.72) | <0.001 |
| Female | 5.60 (5.50, 5.80) | 9.50 (7.20, 11.20) | <0.001 |
| **BMI (z-score)** |  |  |  |
| Total | 2.50 (1.96, 3.05) | 2.47 (1.80, 3.09) | 0.784 |
| Male | 2.96 (2.12, 3.40) | 2.50 (1.89, 3.33) | 0.637 |
| Female | 2.43 (1.82, 2.94) | 2.39 (1.77, 3.00) | 0.991 |
| **ACR (mg/mmol Cr)** |  |  |  |
| Total | 0.20 (0, 0.55) | 0.71 (0.2, 2.7) | <0.001 |
| Male | 0 (0, 0.14) | 0.60 (0.16, 1.71) | 0.001 |
| Female | 0.40 (0.16, 0.78) | 0.84 (0.23, 3.27) | 0.018 |
| **eGFR (ml/min/1.73m<sup>2</sup>)** |  |  |  |
| **Total** | 139.40 (123.60, 153.40) | 143.10 (131.7, 156.07) | 0.063 |
| **Male** | 147.50 (136.6, 163.8) | 154.80 (140.25, 169.98) | 0.685 |
| **Female** | 130.10 (119.58, 144.32) | 139.40 (127.35, 150.85) | 0.049 |
| **Hypertension (N, %)** |  |  |  |
| Total |  |  |  |
| Yes | 18/45, 40.0% | 80/159, 50.3% | 0.298 |
| No | 24/45, 53.3% | 72/159, 45.3% |  |
| N/A | 3/45, 6.7% | 7/159, 4.4% |  |
| Male |  |  |  |
| Yes | 6/15, 40.0% | 24/52, 46.2% | 1.000 |
| No | 7/15, 46.7% | 24/52, 46.2% |  |
| N/A | 2/15, 13.3% | 4/52, 7.6% |  |
| Female |  |  |  |
| Yes | 12/30, 40.0% | 56/107, 52.3% | 0.295 |
| No | 17/30, 56.7% | 48/107, 44.9% |  |
| N/A | 1/30, 3.3% | 3/107, 2.8% |  |

Supporting our findings *in vivo*, patients with T2DM displayed increased serum levels of glucose and lactate, and decreased levels of glutamine, in comparison to the control individuals (Fig.5F**, Table S1**). In agreement with our experimental observations, male adolescents without T2DM displayed significantly increased levels of glucose, lactate, and glutamate, compared to females without T2DM. This increase was also observed in males with T2DM, relative to females with T2DM, without reaching statistical significance. Interaction analyses confirmed the independent contribution of sex and diabetes to the levels of each metabolite. While diabetes was an independent modifier of the circulating levels of glucose and lactate, glutamate levels were exclusively modified by sex (P = 0.031, **Table S2**).

### TCA and glutathione cycle metabolites arising from glutamate are increased in male PTECs and in male individuals

Our data suggested that glutamine and glutamate were involved in major metabolic sex differences. We next examined the expression of key glutamine/glutamate transporters and enzymes. We measured *in vivo* the renal levels of *Slc38a3* (important glutamine importer), *Gls* (glutaminase, which converts glutamine to glutamate), *Glud1* (glutamate dehydrogenase, which converts glutamate to α-KG), and *Oxgr1* (G protein-coupled receptor that mediates signaling induced by α-KG, that is not used up in the TCA cycle) (Fig.S5C). Under non-diabetic conditions, female sex was associated with increased kidney expression of all four genes. In turn, hyperglycemia was linked to a significant increase in the kidney levels of *Slc38a3* and *Glud1* exclusively in diabetic male mice. Diabetes and male sex were also associated with the lowest kidney expression of *Oxgr1* (Fig.S5D). These data suggest a higher conversion of glutamine to glutamate in females, but increased utilization of α-KG in the TCA cycle by diabetic and healthy male kidneys, compared to female kidneys.

In addition to being a carbon source for the TCA cycle, glutamate is also a major substrate of the anti-oxidant glutathione cycle^14^. To determine if glutamate-derived metabolites of the TCA and glutathione cycles were altered between male and female PTECs, we analysed the intracellular metabolites of male and female PTECs exposed to normal glucose (**Table S4A,B**). Supporting our findings in animals and humans, male PTECs displayed a significant increase in the intracellular levels of glutamate, compared to female PTECs (Q<0.05) (Fig.6A). Importantly, male sex was linked to an increase in the intracellular levels of 7 metabolites of the TCA cycle, which represent the ‘pro-oxidant’ metabolism of glutamate. Specifically, male PTECs displayed a significant increase in the intracellular levels of citrate, aspartate, and malate (Q<0.05), and a numerical increase in the intracellular levels of isocitrate, α-KG, succinate, and fumarate, in comparison to female PTECs. Female PTECs showed a marked increase in the intracellular levels of pyruvate (Q<0.001), relative to male PTECs (**Table S4C,** Fig.6A). We also identified sex differences in the intracellular metabolites of the glutathione cycle, the ‘antioxidant arm’ of glutamate metabolism. To exert their antioxidant function, some GSH molecules (the reduced form of glutathione) exit the cycle and neutralize the negative charge of reactive oxygen species, while generating GSSG (oxidized form of glutathione) as by-product^56^. Male PTECs showed significantly higher intracellular levels of GSH and GSSG (Q<0.001), compared to female PTECs. In contrast, female PTECs displayed a significant increase in the GSH/GSSG ratio (Q<0.05), an indicator of adequate redox balance^57^. Female cells also displayed significantly increased levels of glutamyl-cysteine (Q<0.001, precursor of GSH) and cysteinylglycine (Q<0.01, product of GSH). While the cycling of glutathione metabolites seems to be more maintained in female PTECs, a higher efflux of GSH may be occurring in male PTECs, indicating a higher demand for antioxidant mechanisms (**Table S4C,** Fig.6A).

**Figure 6.**
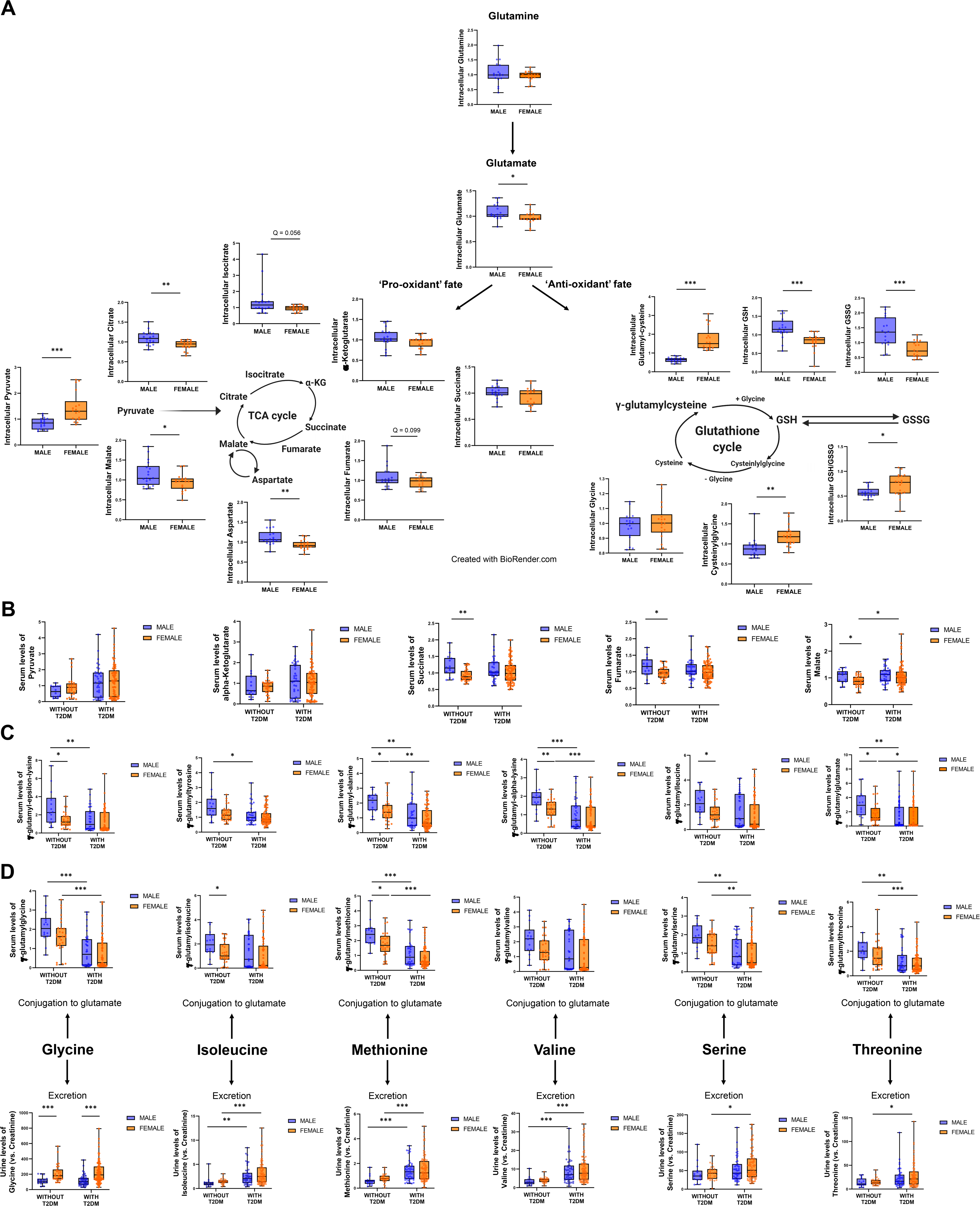
Sex differences in the cell metabolome of PTECs and in the serum metabolome of healthy and diabetic individuals. Metabolome analysis was conducted in male and female PTECs exposed to normal glucose (5.5mM) for 16h (n=3/sex; n=6 replicates/donor). The analysis revealed sex differences in the intracellular levels of key metabolites of the TCA cycle and the glutathione cycle. Differences in intracellular metabolite levels between the two groups were determined using a t-test adjusted for 5% false discovery rate. *q<0.05 (A). Sex differences in the levels of pyruvate, α-KG, succinate, fumarate, and malate were examined in the serum of men without T2DM (n=15), women without T2DM (n=27), men with T2DM (n=47), and women with T2DM (n=91). Data normalization was performed to correct variation emerging from inter-day tuning differences on instrument performance. Each compound was corrected in run-day blocks by registering the medians to 1.00 and normalizing each data point proportionately. Metabolite measurements are thus presented as normalized mass spectrometry-based intensities (B). Male sex was independently associated with increased circulating levels of 9 γ-glutamyl-aminoacids. The levels of 6 of these γ-glutamyl-aminoacids are shown in panel C. Among these individuals, the urinary excretion of glycine, isoleucine, methionine, valine, serine, and threonine was altered by sex and/or diabetes (D). Panels B-E: *p<0.05 and q<0.1; ** p<0.01 and q<0.1; ***p<0.001 and q<0.1. TCA, tricarboxylic acid; α-KG, alpha-ketoglutarate; GSH, reduced glutathione; GSSG, oxidized glutathione; T2DM, type 2 diabetes mellitus. The metabolic cycles in panel A were created with BioRender.com.

Our main findings at the cell metabolome level were reproduced in the iCARE cohort. Females with and without T2DM had numerically increased circulating levels of pyruvate, compared to males. In contrast, male sex was associated with increased circulating levels of TCA cycle metabolites (namely succinate, fumarate, and malate). This increase relative to females was significant only among youth without T2DM, but a similar pattern was observed among patients with T2DM (Fig.6B). We also identified 17 γ-glutamyl-aminoacids, which play a key role as substrates of the glutathione cycle^58^, that were increased in the serum of male individuals (**Table S1,** Figure 6C,D). Among them, circulating levels of γ-glutamyl-epsilon-lysine, γ-glutamyl-alanine, γ-glutamyl-alpha-lysine, γ-glutamylleucine, γ-glutamylglutamate, γ-glutamylisoleucine, and γ-glutamylmethionine were significantly increased in healthy male individuals, compared to females, and the same trend was observed among patients with T2DM. Most of these metabolites (11/17 in males and 13/17 in females) were also significantly decreased by diabetes (**Table S1,** Figure 6C,D). These 17 γ-glutamyl-aminoacids strongly and positively correlated with each other, reinforcing their coregulation (Fig.S6A**, Table S3**). Subsequent interaction analyses showed that sex was an independent modifier of circulating levels of succinate (P = 0.034). Both sex and diabetes were independent modifiers of the levels of γ-glutamyl-α-lysine and γ-glutamylmethionine, while the interaction between the two factors was a modifier of γ-glutamyl-E-lysine levels (**Table S2**). Interestingly, the circulating levels of 15/17 γ-glutamyl-aminoacids significantly and negatively correlated with the levels of glutamine (Q<0.05), while all 17 γ-glutamyl-amino acids significantly and positively correlated with the circulating levels of glutamate. Only 3/17 γ-glutamyl-amino acids correlated with the levels of glucose (Fig.S6B, **Table S3**). These observations suggest that sex differences in the levels of γ-glutamyl-amino acid metabolites in the blood may be due to a sex-specific metabolism of glutamine and glutamate by male and female tissues.

The formation of γ-glutamyl-amino acids is based on the conjugation of a glutamate molecule to itself or to another amino acid, via γ-glutamyl bonds^59,60^. In addition to glutamate, ten amino acids can be a source of carbons for the TCA cycle in the tissues^61^. We postulated that increased kidney conjugation of these amino acids would be reflected in their decreased urine excretion. Urine levels of glycine were significantly increased in females with or without T2DM, as compared to males (P<0.001, Fig.6D). Significant interaction was confirmed between sex and urinary levels of glycine (**Table S2**). Females with or without T2DM also displayed increased urinary excretion of isoleucine, methionine, valine, serine, and threonine, compared to males (Fig.6D). It is noteworthy that the pattern of these 6 amino acids in the urine was a mirror-image of the corresponding γ-glutamyl-amino acid levels in the circulation. A higher demand for amino acids as metabolic substrates in the human male kidney, relative to the female kidney, could explain this effect.

### KDM6A orchestrates metabolic homeostasis in female PTECs

Our next goal was to identify transcriptional regulators of sex-specific metabolic changes in PTECs. We studied genes encoding the proteins upregulated by sex hormones in PTECs^31^. To identify regulators that may underpin the sex chromosome and sex hormone effects, we analyzed 4 publicly available gene datasets of human healthy kidneys (NephroSeq)^62–64^. We identified 196 genes increased in male kidneys and 139 genes increased in female kidneys (Fig.S7, **Table S5**), and used these genes to computationally predict their transcriptional regulators (see Supplemental Methods).

We first focused on the datasets linked to female sex hormones. We identified 49 regulators significantly enriched among the 18 proteins increased by EST, but not by DHT. We also identified 339 regulators significantly enriched exclusively among genes upregulated in the female kidney, but not in the male kidney (**Table S6,** Fig.7A). Among these 339 regulators, we identified the X-linked lysine demethylase KDM6A, also known as UTX, which was increased at transcriptional level in the female kidney (Fig.7B). The main function of KDM6A is to activate gene expression by removing two methyl groups from H3K27me3, a repressive mark on DNA histones^65,66^. KDM6A mediates proliferation and metabolism, and influences glutamine demands, as it requires α-KG to function^67,68^.

**Figure 7.**
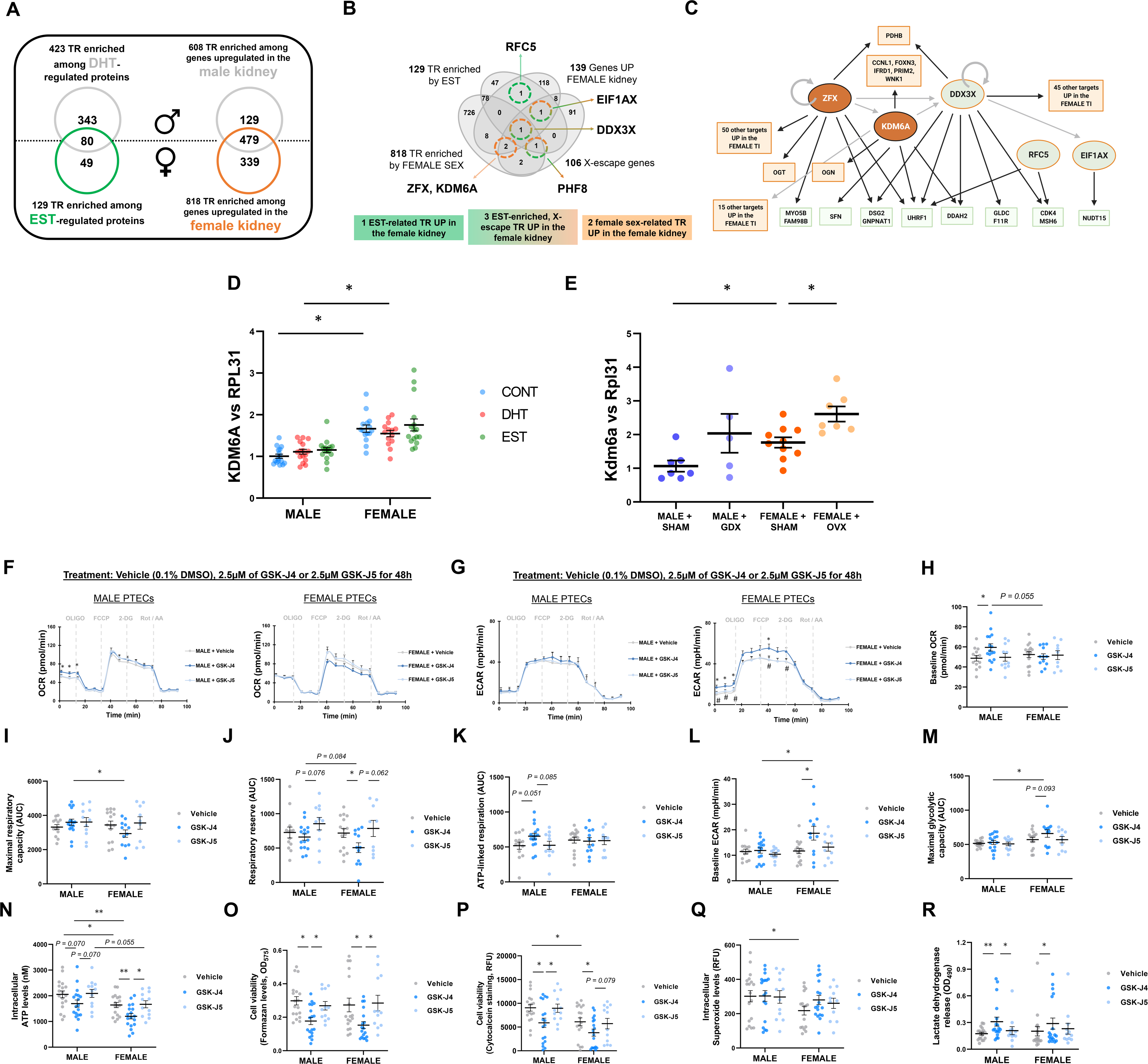
Identification of KDM6A as a regulator of the metabolic function of female PTECs. Kidney genes significantly altered by sex (Nephroseq database), and proteins altered by sex hormones were analyzed using CATRIN transcriptional regulator (TR) database. The analysis revealed a significant enrichment (p<0.05) of 129 TRs among EST-regulated proteins. In turn, 818 TRs were significantly enriched (q<0.05) among genes upregulated in the human female kidney (A). The Venn diagram illustrates the overlap between TRs significantly enriched among kidney molecular signatures linked to male sex, female sex, DHT, and EST. Key TRs linked to female sex are highlighted in orange, while TRs linked to EST are highlighted in green (B). Key TRs and targets emerging from the analysis and relevant to female sex are illustrated (C). The color of each box indicates that the TR/target was enriched/increased by female sex (orange) and/or by EST (green). KDM6A gene expression was determined in male and female PTECs exposed to vehicle (CONT), 100nM DHT, or 100nM EST for 16h, and normalized to RPL31 (n=3/sex; n=4-6/treatment) (D). Kidney *Kdm6a* gene expression was assessed in 19-week-old male and female mice subjected to sham-operation, gonadectomy (GDX) or ovariectomy (OVX) (n=5-10 animals/group) (E). Male and female PTECs were treated with vehicle (DMSO), 2.5μM of GSK-J4 (KDM6A inhibitor), or 2.5μM of GSK-J5 (inactive analog) for 48h (n=3/sex; n=4-8/treatment). After treatment, OCR and glycolysis (ECAR) were measured in a Seahorse XFe96 analyzer (F-G). To induce metabolic stress, the following sequence of drugs was injected: 1μM oligomycin, 0.3μM FCCP, 100mM 2-DG, 1mM Rot/AA. Baseline OCR (H), maximal respiratory capacity (I), reserve respiratory capacity (J), and ATP-linked respiration (K), were calculated from the OCR curves in panel F. In turn, basal glycolysis (L) and maximal glycolytic capacity (M) were calculated from the ECAR curve in panel G. Cell viability was assessed by measuring endogenous formazan formation (MTT assay) (N), uptake of cytocalcein (viability dye) (O), and intracellular ATP levels (P). Intracellular superoxide ion levels (Q) and release of lactate dehydrogenase (R) were also assessed. In panels F-G, a t test was used at each time point to assess statistical significance between the two groups In panels D-E and H-R, Group-to-group differences were determined using pairwise t tests for variables following a normal distribution, and Mann-Whitney tests for variables with a non-parametric distribution. *p<0.05; **p<0.01. PTECs, proximal tubular epithelial cells; CONT, control; DHT, dihydrotestosterone; EST, estradiol; AUC, area under the curve; ECAR, extracellular acidification rate; OCR, oxygen consumption rate; FCCP, p-trifluoromethoxy carbonyl cyanide phenyl hydrazone; 2-DG, 2-deoxyglucose; Rot, rotenone; AA: antimycin A; DMSO, dimethyl sulfoxide; RFU, relative fluorescence units. The illustration in panel C was created with BioRender.com.

Little is known about the role of KDM6A in the kidney, and whether such role is sex-specific. In this study, KDM6A was predicted to regulate 22 genes increased in the female kidney (Q<0.05). Five of these genes (CCNL1, FOXN3, IFRD1, PRIM2, and WNK1) are involved in cell cycle control and proliferation^69^. Together with two additional X-linked regulators (ZFX and DDX3X), KDM6A was predicted to participate in the regulation of the mitochondrial pyruvate dehydrogenase gene PDHB (Fig.7C). KDM6A, ZFX, DDX3X, and PDHB genes were increased in the human female kidney, relative to the male kidney (**Table S5**). KDM6A expression was also significantly increased in female compared to male PTECs (Fig.7D). Female mice had significantly increased kidney gene levels of *Kdm6a* compared to males, even after ovariectomy (removal of estrogens) (Fig.7E).

Given the conserved sex-biased expression of renal KDM6A, and its link to metabolism, we investigated the role of KDM6A in male and female PTECs. Pharmacological inhibition of KDM6A impaired metabolic function and viability of PTECs in a sex-specific fashion (Fig.7F-P). Treatment with the prototypic KDM6A inhibitor GSK-J4^70,71^, but not the inactive control GSK-J5, decreased mitochondrial function (Fig.7F) and increased glycolysis (Fig.7G), in female PTECs. Interestingly, KDM6A inhibition in female PTECs reduced their maximal respiratory capacity (Fig. 7I) and significantly decreased their respiratory reserve (Fig.7J), without affecting ATP-linked respiration (Fig.7K). KDM6A inhibition in female PTECs also resulted in a significantly increased basal glycolytic rate (Fig.7L) and a higher maximal glycolytic capacity (Fig.7M). A metabolic shift toward reduced mitochondrial respiration and increased glycolysis may result in a decreased capacity to generate ATP, compromising proliferation and viability^72^. Increased glycolysis in female PTECs exposed to GSK-J4 was accompanied by a significant decrease in intracellular ATP (Fig.7N) and cell viability (Fig.7O,P), elevated levels of superoxide ion (Fig.7Q) and a significant increase in cell death (Fig.7R). KDM6A may maintain metabolic homeostasis in female PTECs, helping them prioritize the mitochondrial use of energy substrates over glycolysis. In male PTECs, KMD6A inhibition significantly increased basal respiration (Fig.7H), decreased cell viability (Fig.7N-P), and increased cell death (Fig.7R). However, these changes were not linked to altered glycolysis in male PTECs (Fig.7G, L, M).

### HNF4A mediates the deleterious effects of androgens in male PTECs

We next examined transcriptional regulators linked to male sex and sex hormones in the kidney. We identified 343 factors significantly enriched among the 60 proteins upregulated by DHT in male PTECs, but not among the proteins altered by EST. We also identified 608 factors enriched among the genes upregulated in the male kidney (**Table S6,** Fig.8A). We were particularly interested in hepatocyte nuclear factor 4 alpha (HNF4A), predicted to regulate DHT-induced proteins in male PTECs, and genes increased in the human male kidney (Fig.8B). HNF4A mediates fatty acid, glucose and glutamine metabolism^73–77^, and is considered a central regulator of the kidney cell metabolism^78–80^. HNF4A was predicted to regulate the expression of 6 enzymes that were upregulated by DHT in male PTECs^31^, and that play a critical role in the metabolism of glucose (GPI, TKT), glutamine/glutamate (GGH, GLUD1, GLS), or both (GNPNAT1). HNF4A was also predicted to regulate 6 transcription factors encoded on the Y chromosome, either directly (ZFY, EIF1AY, RPS4Y1), or indirectly (UTY, DDX3Y, KDM5D) (Fig.8C). All 6 Y-linked regulators were exclusively expressed at the gene level in male human kidneys and/or male PTECs (**Table S5,** Fig.8B,C, Fig.S8).

**Figure 8.**
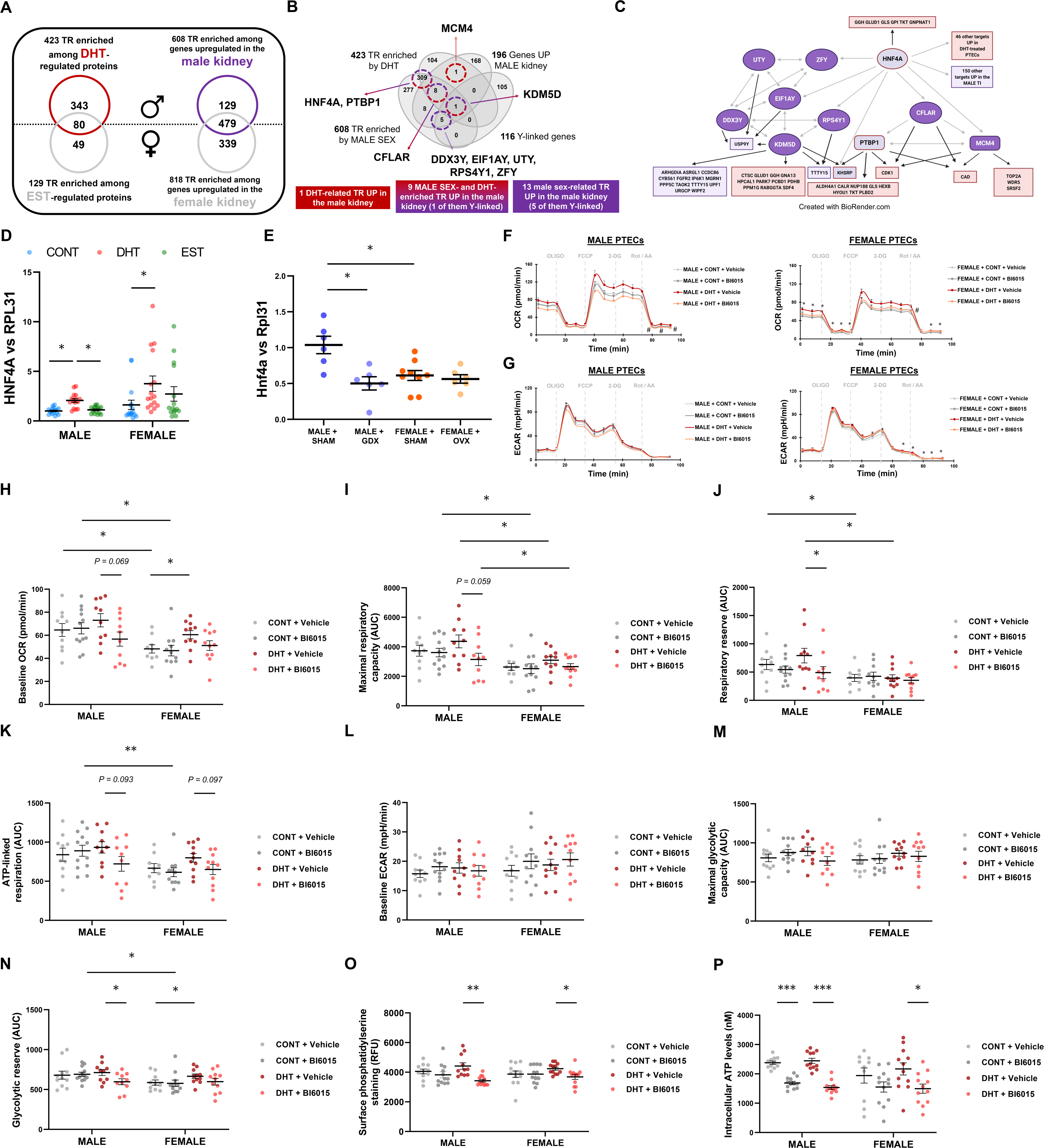
Identification of HNF4A as a mediator of the metabolic effects of DHT in male PTECs. To identify transcriptional regulators (TRs) responsible for the observed metabolic sex differences, kidney genes significantly altered by sex (Nephroseq database) and proteins altered by sex hormones^31^ were analyzed using CATRIN TR database. The analysis revealed a significant enrichment (p<0.05) of 423 TRs among DHT-regulated proteins. In turn, 608 TRs were significantly enriched (q<0.05) among genes upregulated in the human male kidney (A). The Venn diagram illustrates the overlap between TRs significantly enriched among kidney molecular signatures linked to male sex, female sex, DHT, and EST. Key TRs linked to male sex are highlighted in purple, while TRs of interest linked to DHT are highlighted in red (B). Relevant TRs and targets emerging from the analysis and relevant to male sex are illustrated. The color of each box indicates that the TR/target was enriched/increased by male sex (purple) and/or by DHT (red). HNF4A gene expression was determined in male and female PTECs exposed to vehicle (CONT), 100nM DHT, or 100nM EST for 16h, and normalized to RPL31 (n=3/sex; n=4-6/treatment) (D). Kidney *Hnf4a* gene expression was assessed in 19-week-old male and female mice subjected to sham-operation, gonadectomy (GDX) or ovariectomy (OVX) (n=5-10 animals/group) (E). Male and female PTECs were treated for 16h with vehicle or DHT in the presence or absence of DMSO or 0.1μM BI6015 (HNF4A inhibitor) (n=3/sex; n=3-4/treatment). After the treatment, OCR and glycolysis (ECAR) were measured in a Seahorse XFe96 analyzer (F-G). To induce metabolic stress, the following sequence of drugs was injected: 1μM oligomycin, 0.3μM FCCP, 100mM 2-DG, 1mM Rot/AA. Baseline OCR (H), maximal respiratory capacity (I), reserve respiratory capacity (J), and ATP-linked respiration (K), were calculated from the OCR curves in panel F. In turn, basal glycolysis (L), maximal glycolytic capacity (M), and glycolytic reserve (N) were calculated from the ECAR curve in panel G. Surface levels of phosphatidylserine (PS) were measured as a marker of early apoptosis (O). Intracellular ATP levels were also measured (P). Panels F-G: *p<0.05 vs CONT + Vehicle; #p<0.05 vs DHT + Vehicle. Panels D-E, H-P: *p<0.05; **p<0.01. PTECs, proximal tubular epithelial cells; CONT, control; DHT, dihydrotestosterone; EST, estradiol; AUC, area under the curve; ECAR, extracellular acidification rate; OCR, oxygen consumption rate; FCCP, p-trifluoromethoxy carbonyl cyanide phenyl hydrazone; 2-DG, 2-deoxyglucose; Rot, rotenone; AA: antimycin A; RFU, relative fluorescence units. The diagram in panel C was created with BioRender.com.

**Figure 9.**
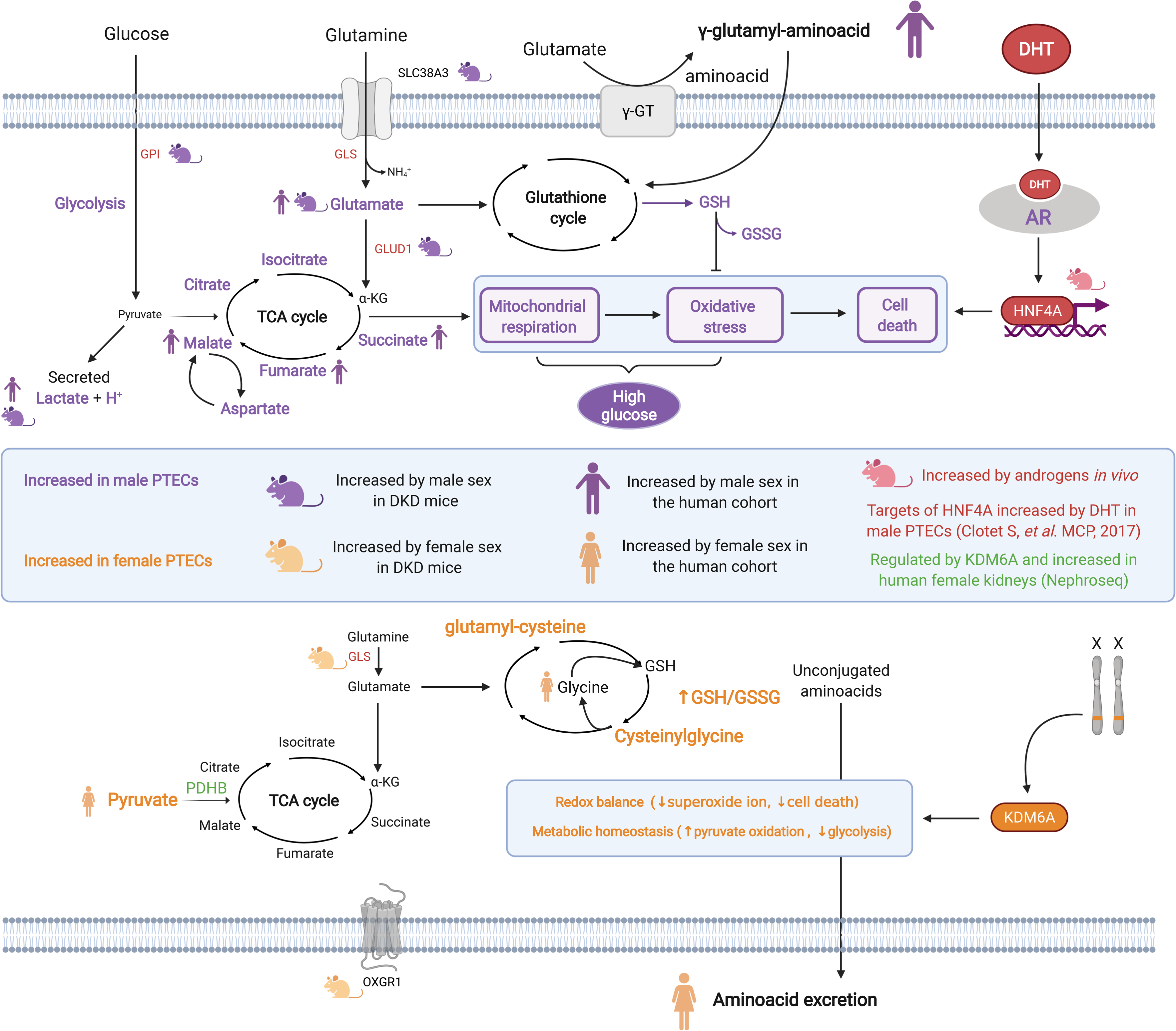
Proposed model based on the integration of *in vitro*, *in vivo,* and human findings. Cell sex and sex hormone effects on kidney metabolism are mediated by KDM6A and HNF4A, respectively, and relate to sex differences in the levels of glucose- and glutamine-derived metabolites. In male PTECs, increased glucose consumption, mitochondrial respiration, oxidative stress, and cell death are enhanced by DHT via AR- and HNF4A-dependent mechanisms. In agreement, metabolic targets of HNF4A involved in glycolysis (GPI) and glutamine anaplerosis (GLS, GLUD1) were upregulated by DHT in male PTECs^31^. Concordantly, increased gene levels of SLC38A3 and GLUD1 pointed towards increased glutamine anaplerosis in diabetic male kidneys. Enhanced renal utilization of glucose and glutamine in males is linked to increased levels of glutamate, malate, fumarate, succinate, GSH, GSSG, and γ-glutamyl-aminoacids, in PTECs and/or in the blood of healthy and diabetic mice and humans, suggesting a higher activation of the TCA cycle and, in consequence, a higher demand for antioxidant mechanisms. In female PTECs, metabolic homeostasis is regulated by KDM6A-dependent mechanisms, and linked to increased cellular levels of pyruvate, glutamyl-cysteine, and cysteinylglycine, and to a higher GSH/GSSG ratio, suggestive of a favorable redox balance. Increased aminoacid excretion in non-diabetic and diabetic female individuals suggest a lower demand for aminoacid utilization in the glutathione cycle of female tissues.

In the kidney, HNF4A has been proposed as the main partner of AR^45^. Along the same lines, DHT treatment significantly increased HNF4A gene expression in both male and female PTECs (Fig.8D). Similarly, kidney *Hnf4a* gene expression was significantly increased in male compared to female mice, and this increase was completely prevented in gonadectomized males (Fig.8E). In contrast, HNF4A gene expression did not differ between human male and female healthy kidneys, in publicly available datasets (**Table S5**). In summary, HNF4A was modified by male sex hormones, but not sex. We wondered if HNF4A also mediated the metabolic effects of DHT. We thus inhibited HNF4A in DHT-treated male and female PTECs, using BI6015, a pharmacological antagonist of HNF4A^81^. Co-treatment with BI6015 significantly prevented the DHT-induced increase in mitochondrial respiration, glycolysis, and apoptosis, and this effect was more pronounced in male PTECs (Fig.8F-O). HNF4A inhibition decreased basal respiration (Fig.8H), maximal respiratory capacity (Fig.8I), respiratory reserve (Fig.8J), ATP-linked respiration (Fig.8K), maximal glycolytic capacity (Fig.8M), and glycolytic reserve (Fig.8N) of DHT-treated male PTECs. These metabolic effects were associated with a decrease in apoptosis (Fig.8O). BI6015 also reduced these parameters in female PTECs exposed to DHT, but in a more subtle fashion (Fig.8H-O). Interestingly, HNF4A inhibition significantly reduced the cellular ATP levels in DHT-treated PTECs, but also in control-treated cells, suggesting that HNF4A also governs ATP-producing mechanisms in PTECs independently from androgens (Fig.8P). These findings suggest that HNF4A mediates the deleterious metabolic effects of androgens, but also has androgen-independent metabolic effects, in male PTECs.

## DISCUSSION

The existence and clinical implications of the sex-based progression of DKD are well recognized^82–85^. Despite its importance, the underlying causes of the sex dimorphism in health or in disease remain unknown. Here, we demonstrate that: 1) male sex and DHT promote oxidative metabolism and cell injury in PTECs, while female sex promotes a more quiescent metabolism; 2) male PTECs are more susceptible to high glucose-induced injury and display a higher mitochondrial utilization of glucose and glutamine, while female PTECs show a predilection for pyruvate; 3) male sex is linked to increased levels of lactate, glutamate, TCA cycle, and glutathione cycle metabolites, while female sex is linked to increased levels of pyruvate, *in vitro*, *in vivo* and in healthy and diabetic individuals; and 4) KDM6A orchestrates metabolic homeostasis in female PTECs, while HNF4A mediates the deleterious effects of androgens in male PTECs. This study supports our previous proteomics findings^31^, and identifies, for the first time, a compelling link between glucose/glutamine metabolism and sex, that may explain the roots of sex dimorphism in DKD and possibly other kidney diseases.

Growing evidence demonstrates that the contribution of sex chromosomes in health and disease is broader than previously thought^86^. Genes escaping from the inactivation of one of the X chromosomes can be elevated in adult female tissues^65^, while Y chromosome genes are expressed by adult, somatic male tissues^87,88^. In the kidney, genes encoded by sex chromosomes can exhibit sex-biased expression and exert important functions^89–91^. In mammals, both sex chromosomes and sex hormones control tissue morphogenesis, organ size and metabolism, at different stages of life^92,93^. These effects are evolutionary connected to the specific metabolic demands of male and female tissues^86^, which may result in a different susceptibility to metabolic diseases^92,94^. In DKD, sex differences in progression have been linked to a differential activation of mechanisms controlling renal hemodynamics, such as the renin-angiotensin system^32,33^ or nitric oxide signaling^33,95,96^. We previously uncovered a link between androgens and increased expression of metabolic enzymes in PTECs, and validated our key findings in two models of diabetes^31^. We now considered the sex of the kidney cell as a principal study variable. Our novel approach led to the identification of fundamental sex differences in the metabolism of kidney cells, incentivising a paradigm shift in the study of sex differences in kidney health and disease.

Our main observation at the functional level is that male sex and DHT induce a more energetic and oxidative metabolic phenotype in PTECs, based on increased glucose consumption, glycolysis, mitochondrial respiration, oxidative stress, and apoptosis. Supporting these findings, we and others have shown increased oxidative stress levels in the kidneys of non-diabetic and diabetic male compared to female animals^31,97,98^. Increased ROS production by male mitochondria has also been described in other tissues^99–101^. We have now demonstrated that this sex dimorphism is conserved in PTECs, and attributed to both cell sex and sex hormones. PTECs are the most abundant and energy-demanding cell type in the kidney and possess a high mitochondrial content^102^. Mitochondrial respiration is the main ATP-producing mechanism, but also results in the generation of oxidative stress^103^. One may postulate that the higher rates of mitochondrial respiration in male PTECs explain the augmented levels of oxidative stress in male kidneys. Therefore, identifying and targeting the metabolic mechanisms responsible for this mitochondrial hyperactivity may help diminish oxidative stress and DKD progression in males.

Increased mitochondrial utilization of glucose and glutamine by male PTECs may explain their increased rates of respiration. Although fatty acid oxidation (FAO) is considered the main energy-producing mechanism in proximal tubules^104^, mitochondrial glucose and glutamine utilization are also crucial for the kidney cell homeostasis^105–107^. In DKD, defective renal FAO is accompanied by increased glycolysis and altered glutamine metabolism^108–110^. A higher influx of glucose to kidney cells may compromise the balance between the mitochondrial and glycolytic fates of glucose, resulting in enhanced lactate secretion. This shift leads to increased demands for glutamine to maintain acid/base homeostasis and to sustain the TCA cycle, through conversion to glutamate^15,20,47^. The increase in glucose and glutamine utilization by the mitochondria of male PTECs may explain why male kidneys are more prone to DKD injury than female kidneys. Indeed, male PTECs exposed to high glucose showed an initial increase in glycolysis and oxygen consumption, which remained higher than in female cells. These changes were followed by a more rapid decline ATP levels, a sign of injury in kidney cells exposed to high glucose^111,112^. Increased glycolysis and enhanced mitochondrial utilization of glutamine in male PTECs could predispose them to a more severe injury under high-glucose conditions, when glycolysis is favoured^113,114^.

Our approach revealed sex differences in glutamate-derived metabolites in PTECs that were conserved in a relevant clinical setting. Male PTECs showed increased levels of glutamate and 7 metabolites of the TCA cycle, the metabolic pillar of mitochondrial respiration^115^. Of those, glutamate, malate, succinate and fumarate were also increased in the blood metabolome of male subjects with and without T2DM. Krumsiek *et al*. identified increased levels of circulating malate in healthy males, compared to females^116^. *In vivo*, succinate levels were increased in the kidneys of male mice, compared to females, and changed in a sex-specific fashion in the setting of T1DM^117^. Although we did not measure the TCA cycle metabolites in the urine, a previous analysis of the healthy human urine metabolome showed that males had a lower excretion of malate, citrate, succinate, and fumarate than females^118^. Our findings point towards a higher activation of the TCA cycle in the male kidney, which could explain the reduced excretion of TCA metabolites in males^118^. Our sex-based observations regarding the TCA cycle may be important in DKD^119–123^. For example, altered urine and plasma levels of glutamate, citrate, malate, and fumarate predicted CKD progression in T2DM, especially in male patients^119,120,123^. Increased TCA cycle metabolites in males may reflect hyperactivation of this pathway and impaired redox balance in kidney cells.

Male sex was also linked to increased cellular levels of GSH and GSSG, the effector metabolites of the glutathione cycle^56^. In turn, female PTECs displayed elevated levels of glutamyl-cysteine and cysteinylglycine as well as elevated GSH/GSSG ratio (indicative of a favourable redox balance^57^). Supporting our findings, decreased oxidative stress in female tissues has been linked to higher antioxidant capacity, increased GSH/GSSG ratio, and differential activation of glutathione-related genes and enzymes under healthy^99,100,124^ and diabetic conditions^125,126^. A higher efflux of GSH may be occurring in male PTECs, since they displayed higher oxidative stress than female PTECs and may have a higher demand for antioxidant mechanisms. In the human cohort, we identified 17 γ-glutamyl-aminoacids intimately linked to the glutathione cycle that were significantly increased by male sex and decreased by diabetes. Similarly, Krumsiek *et al.* found higher levels γ-glutamylleucine, γ-glutamylvaline, γ-glutamylphenylalanine, γ-glutamylisoleucine, γ-glutamyltyrosine in male compared to female sera^116^. Our work confirms the link between sex and circulating γ-glutamyl-aminoacids and demonstrates that this aspect of sex dimorphism is conserved in the setting of T2DM. In urine, we found a strong association between female sex and increased excretion of glycine. This association was observed before in healthy subjects^118^, and we now describe it in T2DM. The urine excretions of isoleucine, methionine, valine, serine, and threonine were increased by female sex and also by diabetes, mirroring the pattern of their corresponding γ-glutamyl-aminoacids in the circulation. While glycine is a natural precursor of GSH, all 6 aminoacids can conjugate to glutamate and facilitate its utilization in the glutathione cycle^127,128^. A higher demand for these aminoacids as substrates of the glutathione cycle in the male kidney, relative to the female kidney, could explain their increased excretion in females.

The different type of diabetes between the animals (T1DM) and humans (T2DM) studied here may represent a limitation. However, renal metabolic alterations in DKD are triggered by hyperglycemia and have been described in both T1DM and T2DM^15,129^. Furthermore, Akita mice show microalbuminuria and hyperfiltration^53,130^, resembling the renal function profile of the iCARE participants with T2DM. In contrast to prior metabolomics studies^119–123^, we identified metabolite changes that occurred early after the onset of diabetes, and before the establishment of overt DKD. The value of reporter metabolites that can infer alterations in tissue homeostasis in a non-invasive fashion, and at early stages of disease, is increasingly recognized in nephrology^131–133^.

At the mechanistic level, we identified a link between KDM6A and female kidney metabolism. KDM6A mediates epigenetic modifications^67^, which are linked to metabolism and “molecular memory” in DKD^20,134^. KDM6A can escape from X inactivation, resulting in increased expression due to a double gene dosage in female (XX), compared to male tissues (XY)^65,125,135,136,124^. Here we show increased KDM6A expression in female PTECs and mouse kidneys. KDM6A inhibition in female PTECs led to a metabolic shift characterized by increased glycolysis, oxidative stress, and apoptosis, suggesting that KDM6A contributes to maintenance of metabolic homeostasis. In agreement, studies in autoimmunity models revealed that KDM6A mediates metabolism and exerts sex-specific functions in macrophages and T cells^68,136^. Furthermore, female chromosomes and increased KDM6A exerted protective actions in mice with cardiac ischemia^137^. Our female PTECs showed higher levels of pyruvate than male PTECs. In turn, our bioinformatics analyses revealed that KDM6A participates in the regulation of PDHB. This enzyme catalyzes the mitochondrial use of pyruvate, preventing its conversion to lactate through glycolysis^138^. By lowering glycolysis, KDM6A may contribute to an increased cytosolic pool of pyruvate in female PTECs. This idea is supported by data generated in immune cells^136,139,140^. New lines of investigation of sex differences and epigenetics may help identify important sex-based regulatory mechanisms in kidney health and disease.

Finally, we have identified a link between androgens and HNF4A, a central regulator of glucose and glutamine metabolism^73–77,141^. In the kidney, HNF4A is highly expressed in PTECs^142,143^ and has been proposed as the main co-regulator of AR^144^. Accordingly, androgens increased HNF4A levels in our PTECs, while orchidectomy decreased kidney *Hnf4a*. Our findings reinforce the link between sex hormones and HNF4A in the kidney. We have also demonstrated that the deleterious metabolic effects of androgens in male PTECs are mediated by HNF4A. At first glance, these findings may seem contradictory, as previous studies suggest that the renal metabolic actions of HNF4A are beneficial^145,146^. HNF4A also regulates gene expression via androgen-independent mechanisms^143,145,147,148^. While androgen-independent HNF4A actions seem essential for PTECs, androgen-dependent mechanisms of this transcription factor may result in mitochondrial hyperactivity and predispose to oxidative stress. In diabetes, high glucose alters the levels and activity of HNF4A^149,150^. In turn, HNF4A regulates genes altered in PTECs from diabetic patients^151^. Since both DHT and HNF4A control metabolic processes altered by diabetes, they may play a synergistic role in DKD^31,73,152,153^.

Our work sheds new light on the sex-based dimorphism in kidney metabolism and DKD. Our sex-balanced approach spearheads new research paradigms through the investigation of the distinct metabolic wiring of male and female kidney cells that can contribute to homeostasis and disease. We are the first to identify fundamental sex differences in the kidney cell metabolism of glucose and glutamine, linking them to early changes in glutamate-derived metabolites under diabetic and non-diabetic conditions, and to sex-specific mechanisms of transcriptional and metabolic regulation. Our findings pave the way to new avenues of research based on patient sex, with the potential to improve monitoring and prevention of DKD in men and women.

## METHODS

### Cell culture

PTECs were purchased from Lonza Walkersville Inc (Walkersville, MD). They were cultured in custom-made Dulbecco’s modified Eagle’s medium (DMEM) containing 5.55mM D-glucose, 4mM L-glutamine, and 1mM sodium pyruvate. Growth DMEM media was supplemented with 10% v/v dialyzed fetal bovine serum (FBS), 10ng/mL EGF, 1x of Transferrin/Insulin/Selenium (Invitrogen), 0.05M hydrocortisone, 50units/mL penicillin, and 50g/mL streptomycin, as previously described^31,154^. All experiments were performed at passage 5. To study the effects of cell sex, PTECs from 4 different male donors and 5 different female donors were studied. To study the effects of sex hormones in male and female PTECs, cells were serum-starved for 24h and treated with 100nM DHT (D-073 Sigma) or 100nM EST (3301 Sigma) for 16h or 24h. Ethanol-treated cells were used as controls (CONT). After stimulation, cell media were collected and centrifuged at 2000G for 10min at 4°C, and supernatants were stored at -80°C. For gene expression experiments, cells were washed with PBS, harvested with trypsin, and snap-frozen at 80°C until further analysis. See Supplemental Methods for experimental details on high glucose and pharmacologic inhibition studies.

### Assessment of metabolic function in kidney cells

Glycolysis was assessed in male and female PTECs by measuring the extracellular acidification rate (ECAR) in a Seahorse XFe96 analyzer (Agilent). Oxygen consumption rate (OCR) was also monitored. Upon confluence, cells were detached with 0.25% trypsin for 5min at 37°C, and subsequently seeded in a Seahorse XFe96 Cell Culture Microplate at a density of 15,000 cells/well in 100µL of DMEM complete media. After letting them adhere for 4-6h, PTECs were starved for 24h and exposed to the treatment of interest. One hour prior to the assay, starvation DMEM media was removed, and cells were washed with phenol-free basal media (Agilent) and exposed to 150µL of minimal substrate assay media. The assay media was prepared freshly by adding 2mM glutamine, 1mM pyruvate, and 5.55mM glucose (normal glucose experiments) or 25mM glucose (high glucose experiments) to the basal media. For substrate restriction experiments, the assay media was prepared with only glucose (5.55mM), glutamine (2mM), or pyruvate (1mM). The same concentrations of sex hormones (100nM for normal glucose conditions and 1nM for high glucose conditions) were maintained during this acclimatization step. See Supplemental Methods for details about induction of metabolic stress and calculation of ECAR- and OCR-related parameters in PTECs.

### Glucose uptake

Glucose uptake in male and female PTECs was measured with a Glucose Uptake-Glo™ Assay Kit (Promega), following manufacturer instructions (see Supplemental Methods).

### Cell injury

#### Cellular oxidative stress and apoptosis

Oxidative stress in male and female PTECs was assessed by measuring the intracellular levels of superoxide ion with the Cellular ROS Assay Kit (Red) (Abcam) following the manufacturer instructions. In turn, early apoptosis was assessed by measuring the levels of phosphatidylserine on the cell surface using the Apoptosis/Necrosis Assay Kit (Abcam) (see Supplemental Methods).

#### Lactate dehydrogenase release

Release of LDH in the cell supernatant was assessed as a marker of cell death in male and female PTECs using the CytoTox96® Cytotoxicity Assay (Promega) following the manufacturer instructions (see Supplemental Methods).

### Cell viability

#### Cellular ATP levels

Intracellular levels of ATP in male and female PTECs were measured with a CellTiter-Glo 2.0 Assay Kit (Promega), following manufacturer instructions (see Supplemental Methods).

#### MTT assay

Cell viability was also assessed in male and female PTECs using the MTT assay (Sigma), following manufacturer instructions (see Supplemental Methods).

### Cell metabolome

#### Sample preparation

Male and female PTECs were grown on 6-well plates and serum starved as described above. The steady-state intracellular metabolome was then determined after 16h of exposure to normal glucose starvation conditions, using liquid chromatography-mass spectrometry (see Supplemental Methods).

### Animal studies

To assess sex differences in glucose and glutamine metabolism *in vivo*, the kidneys and the plasma of 8-week- and 16-week-old C57BL/6 healthy male and female mice were studied. Male and female diabetic Akita (Ins2WT/C96Y) mice of the same age groups were studied in parallel to assess the effect of sex in the context of type 1 diabetes. To evaluate the role of male and female sex hormones in the kidney levels of key transcriptional regulators, 10-week-old C57BL/6 healthy male and female mice were subjected to gonadectomy (GDX) and ovariectomy (OVX), respectively (see Supplemental Methods).

To assess the effect of androgens on the urine excretion of glucose, lactate, glutamine, and glutamate, we studied the effect of GDX in a different colony of C57BL/6 10-week-old healthy male mice, as previously described ^34^ (see Supplemental Methods).

At the end of each follow-up, morning spot urine was collected through abdominal massage, cleared by centrifugation at 8,000G for 10min, and stored at -80°C until further analysis. Mice were then anesthetized with isoflurane and sacrificed by terminal surgery. Blood was collected by cardiac puncture in heparinized tubes (Sarstedt). Plasma was subsequently obtained by centrifugation at 8,000G for 10min and stored at -80°C. Kidneys were removed, weighted, snap frozen in liquid nitrogen and kept at -80°C until further analysis. Five to eight animals were studied in each experimental group.

Mouse studies were performed at the Division of Comparative Medicine at University of Toronto (healthy controls and diabetic Akita mice), at the Laval University Animal Supply Facility (GDX and OVX male and female mice), and at the Animal Facility of Barcelona Biomedical Research Park (female, male, and GDX male mice). Mice were housed in ventilated cages with full access to chow and water, in a controlled-temperature environment maintained under a 12h light/dark cycle. All experiments were conducted under the guidelines of the University of Toronto Animal Care Committee, the Laval University Animal Care Committee, and the Ethical Committee of Animal Experimentation of Barcelona Biomedical Research Park (CEEA-PRBB). All efforts were made to use the minimal number of mice and minimize animal suffering.

### Targeted metabolite measurements

Levels of glucose, lactate, glutamine, and glutamate were measured in cell supernatants, plasma and urine using a BioProfile® FLEX2™ Automated Cell Culture Analyzer (Nova Biomedical) (see Supplemental Methods).

### Gene expression

Gene expression studies were conducted on RNA extracted from PTEC cell pellets or from 30-50mg of mouse kidney cortex using the RNAeasy Mini Kit (Qiagen) (see Supplemental Methods). After quantifying RNA concentration in a Nanodrop instrument (Thermo) 300-700ng of RNA were retrotranscribed to cDNA using the High-Capacity cDNA Reverse Transcription Kit (Applied Biosystems). For the *in vitro* experiments, male and female PTECs were treated with vehicle, 100nM DHT, or 100nM EST for 16-24h under normoglycemic conditions. In these cells, gene levels of AR, KDM6A, ZFX, DDX3X, HNF4A, ZFY, EIF1AY, RPS4Y1, UTY, DDX3Y, and KDM5D were measured by real-time quantitative PCR using a Power SYBR® Green PCR Master Mix reagent (Applied Biosystems) and normalized to HPRT1 or RPL31. The fluorescent signal was measured in a StepOnePlus System (Applied Biosystems) for 96-well plates, and in a LightCycler® 480 Instrument II (Roche) for 384-well plates. For the *in vivo* experiments, gene levels of *Slc38a3*, *Gls*, *Glud1*, *Oxgr1, Kdm6a, Hnf4a, Zfy, Eif1ay, Uty, and Ddx3y* were measured and normalized to *Hprt1* or *Rpl31*. All primer sequences employed in this study are summarized in **Table S7**.

### Human studies

#### Clinical cohort

We examined sex differences in the blood metabolome of individuals from the iCARE cohort (University of Manitoba). iCARE is a prospective, observational cohort study to identify risk factors for early onset albuminuria and progression of CKD in male and female adolescents with T2DM (n=322)^155^. These patients were studied together with controls without diabetes of the same age group and ethnicity (n=139). Metabolome analysis was performed on serum and/or urine samples collected from a subgroup of 204 participants of the iCARE cohort (15 males without T2DM, 30 females without T2DM, 52 males with T2DM, and 107 females with T2DM). Specifically, metabolite measurements were performed on serum samples from 155/204 cases (14 males and 28 females without diabetes, 36 males and 77 females with T2DM) and on timed overnight/first morning urine samples from 180/204 cases (15 males and 27 females without diabetes, 47 males and 91 females with T2DM) (Fig.S9).

#### Serum and urine metabolome analysis

The 155 serum and 180 urine samples of the iCARE cohort were prepared using the automated MicroLab STAR® system (Hamilton Company). The metabolome was measured in the serum and urine sample aliquots following an untargeted UPLC-MS/MS approach (Metabolon Inc, Durham, North Carolina). Raw data files from the MS runs were extracted, peak-identified, and processed for QC using Metabolon’s hardware and software. Curation and QC processes were conducted by Metabolon data analysts to ensure accurate and consistent identification of true chemical entities. After curation, peak quantification for each metabolite was preformed using the area-under-the-curve (see Supplemental Methods).

### Bioinformatics

#### Determination of sex-biased kidney genes in the human kidney tubulointerstitium

To identify genes differentially expressed between male and female kidneys that could relate to our findings in tubular cells we analyzed publicly available gene expression data in Nephroseq database. The analysis led to the identification of 196 genes significantly upregulated in the male tubulointerstitium and 139 genes significantly upregulated in the female tubulointerstitium (**Table S5,** Fig.S7) (see Supplemental Methods). These two lists of genes were used to identify transcriptional regulators linked to male or female sex, as described below.

#### Prediction of sex-specific transcriptional regulators of human kidney signatures

To identify transcriptional mechanisms linked to the effect of sex hormones in the kidney tubule, we studied computationally which regulators are predicted to target the 60 proteins upregulated by DHT, as well as the 18 upregulated by EST, in PTECs^31^. To account for sex chromosome effects on the kidney tubule, we studied which regulators are predicted to target the 196 genes significantly upregulated in the male tubulointerstitium and the 139 genes significantly upregulated in the female tubulointerstitium (**Table S5,** Fig.S7). We queried these 4 lists of proteins and genes in CATRIN 1.0, a transcriptional regulator database that integrates the findings of 15 stand-alone transcriptional regulator databases (http://ophid.utoronto.ca/Catrin). Regulators of the genes of interest were identified using CATRIN interaction (excluding TF2DNA.experimental) and hypergeometric test was performed to identify which molecules significantly regulated the lists of genes of interest (see Supplemental methods).

### Statistical analysis

The statistical analysis of our *in vitro* and *in vivo* data was performed using GraphPad Prism v9.1.2. Normalized serum and urine metabolome data was analyzed using R^156^ version 4.0.2. Data from 65 metabolites of interest in serum and 19 metabolites in urine was analyzed using three statistical methods (see Supplemental Methods).

## AUTHOR CONTRIBUTIONS

SC-F and AK conceived the study.

SC-F, OZ, HR, and AK participated in study design.

SC-F and OZ carried out the *in vitro* experiments.

SC-F, MR, MJS, AI, JL-P, SP, KC, DS, and JWS retrieved and provided the biospecimens for the *in vivo* studies.

ABD, BW, JMM, and TDB provided clinical and metabolome data from the iCARE cohort for the human studies and helped with the analyses of the clinical data and the conceptual advance.

SC-F, OZ, CP, MK, CMM, HR, and AK analyzed the data.

SF, AS, AB, and MC provided technical support.

SC-F, OZ, CP, MK, IJ, MW, ABD, BW, JMM, TDB, and AK drafted, revised, and edited the paper; and all authors approved the final version of the manuscript.

## ACKNOWLEDGEMENTS

We thank Madhurangi Arambewela, Shilpa Balaji, and Stefan Petrovic for their technical support, and Brenden Dufault for conducting statistical analyses. We would like to thank the patients and their families for participating in the iCARE research study and for generously donating the biospecimens.

---

**Figure S1.**
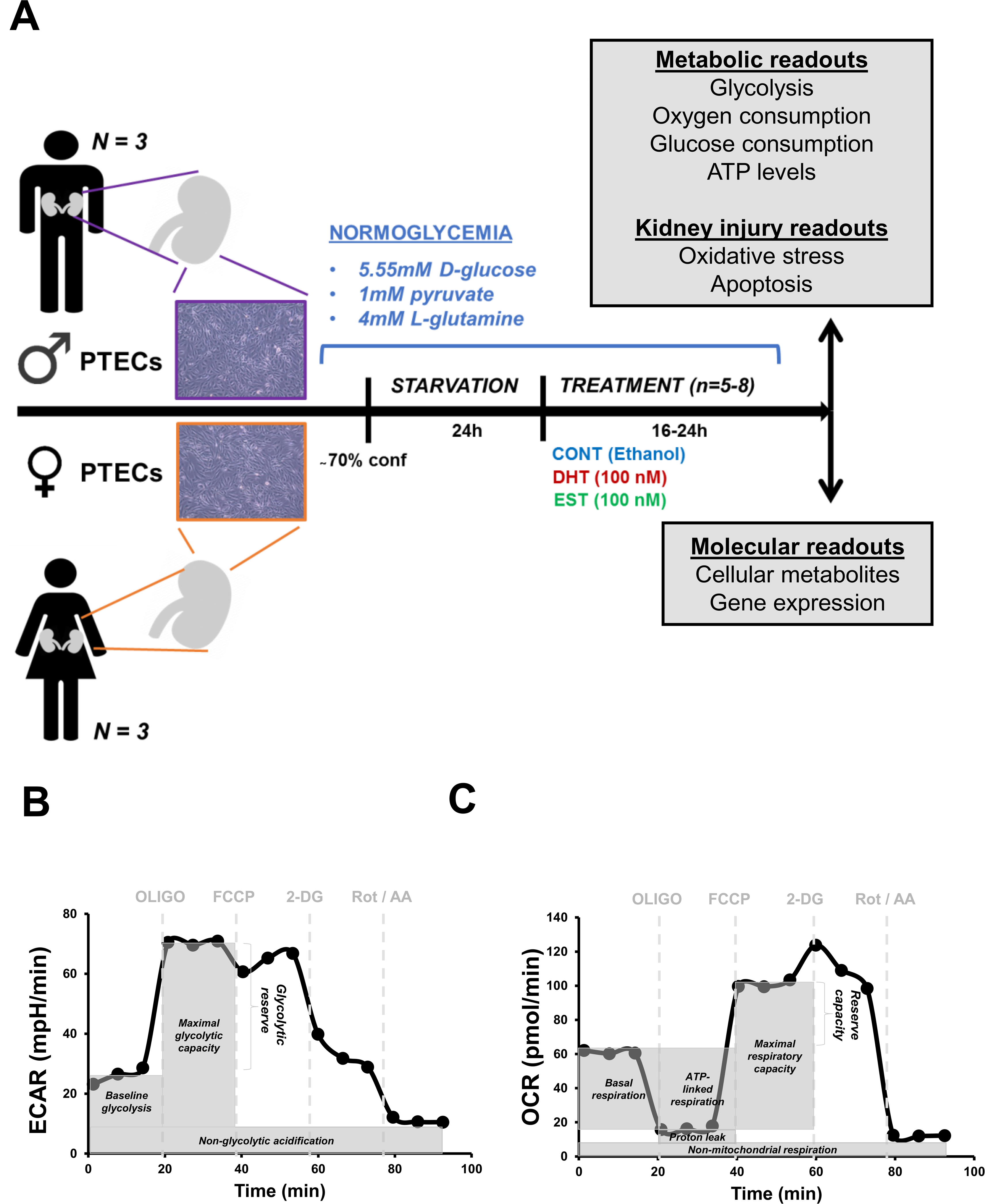
Sex-balanced experimental design. To study the role of cell sex and sex hormones in the setting of normal glucose, PTECs from 4 male and 5 female donors were studied in parallel. The commercial availability of male and female PTECs (Lonza) enabled the simultaneous study of cells from a maximum of 3 donors of each sex. After being serum-starved for 24h, PTECs were exposed to vehicle (CONT), 100nM DHT, or 100nM EST for 16-24h. A series of metabolic, kidney injury, and molecular readouts were measured after treatment (A). As part of the metabolic function studies, baseline glycolysis, maximal glycolytic capacity, glycolytic reserve, and non-glycolytic acidification were assessed by calculating the estimated AUC on different sections of each ECAR curve (B). Basal respiration, ATP-linked respiration, maximal respiratory capacity, reserve capacity, proton leak, and non-mitochondrial respiration were assessed by calculating the estimated AUC on different sections of each OCR curve (C). PTECs, proximal tubular epithelial cells; AUC, area under the curve; DHT, dihydrotestosterone; EST, estradiol; ECAR, extracellular acidification rate; OCR, oxygen consumption rate; OLIGO, oligomycin; FCCP, p-trifluoromethoxy carbonyl cyanide phenyl hydrazone; 2-DG, 2-deoxyglucose; Rot, rotenone; AA: antimycin A.

**Figure S2.**
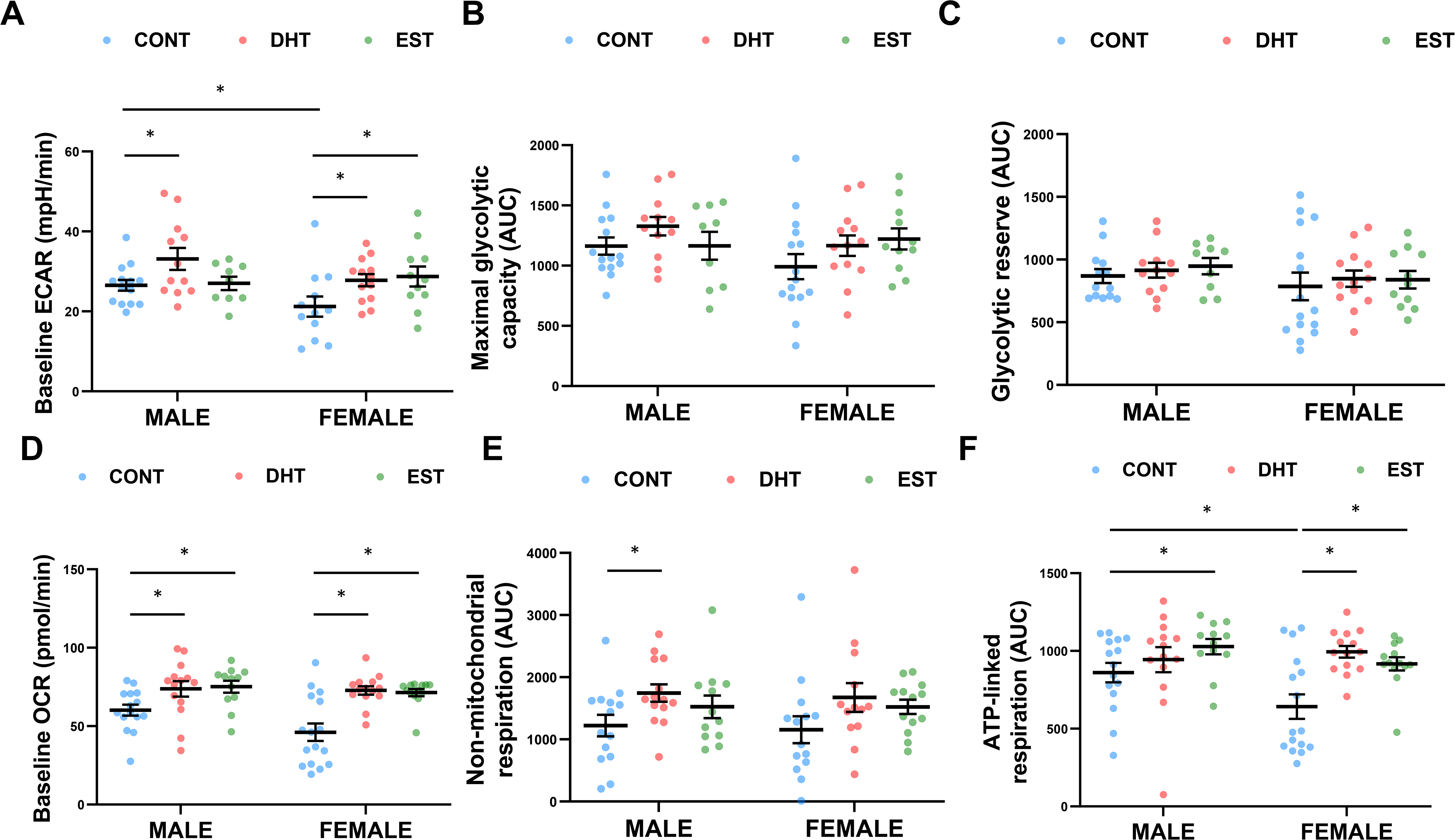
Effect of sex hormones on the metabolism and proliferation of male and female PTECs. Male and female PTECs were exposed to ethanol (CONT), 100nM DHT or 100nM EST for 16-24h (n=3/sex; n=4-6/treatment). Baseline ECAR (A), maximal glycolytic capacity (B), glycolytic reserve (C), basal respiration (D), non-mitochondrial respiration (E), and ATP-linked respiration (F) were calculated from the ECAR and OCR curves in Figure 2A. Group-to-group differences were determined using pairwise t tests for variables following a normal distribution, and Mann-Whitney tests for variables with a non-parametric distribution. *p<0.05. PTECs, proximal tubular epithelial cells; AUC, area under the curve; DHT, dihydrotestosterone; EST, estradiol; ECAR, extracellular acidification rate; OCR, oxygen consumption rate.

**Figure S3.**
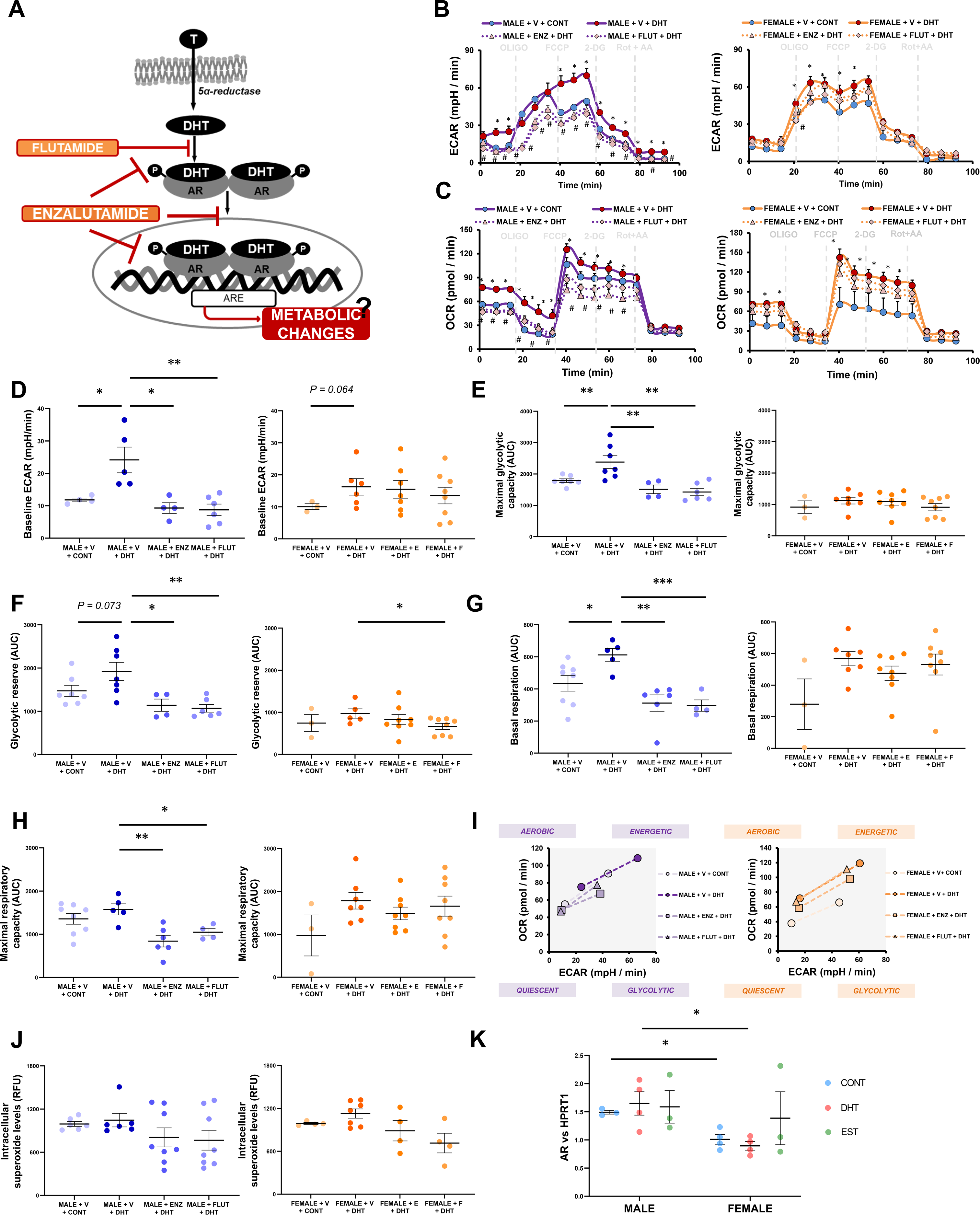
Effect of androgen receptor inhibition on the metabolic function of male and female PTECs. Male and female PTECs were exposed to anti-androgen treatment for 2h prior to stimulation vehicle (CONT) or dihydrotestosterone (DHT) for 16h (n=2/sex; n=4-6/treatment). Two different AR inhibitors were employed: flutamide (FLUT) and enzalutamide (ENZ). FLUT is a selective antagonist of the AR, and competes with androgens like DHT for binding to AR. In turn, ENZ has a novel mode of action targeting AR signalling at three key stages: 1) inhibits androgen binding to AR; 2) inhibits nuclear translocation of the hormone/AR complex; and 3) inhibits binding of the hormone/AR complex to DNA (A). Pre-treatment with either FLUT (1µM) or ENZ (1µM) significantly prevented the DHT-induced increase in glycolysis (B) and mitochondrial respiration (C) in male PTECs and, to a lesser extent, in female PTECs. Accordingly, both FLUT and ENZ significantly reduced baseline ECAR (D), maximal glycolytic capacity (E), in male PTECs. Overall, FLUT and ENZ prevented the change towards a more energetic metabolic phenotype induced by DHT in male PTECs. Although to a lesser extent, this effect was also observed in female PTECs (I). In addition, pre-treatment with FLUT and ENZ reduced the intracellular levels of superoxide ion in DHT-treated male and female PTECs (J). In line with their increased susceptibility to DHT treatment and AR inhibition, male PTECs displayed a significant increase in the gene expression of AR, compared to female PTECs (K). Group-to-group differences were determined using pairwise t tests for variables following a normal distribution, and Mann-Whitney tests for variables with a non-parametric distribution. *p<0.05; **p<0.01; ***p<0.001. PTECs, proximal tubular epithelial cells; AUC, area under the curve; DHT, dihydrotestosterone; EST, estradiol; ECAR, extracellular acidification rate; OCR, oxygen consumption rate; AR, androgen receptor; HPRT1, hypoxanthine-guanine phosphoribosyltransferase; RFU, relative fluorescence units.

**Figure S4.**
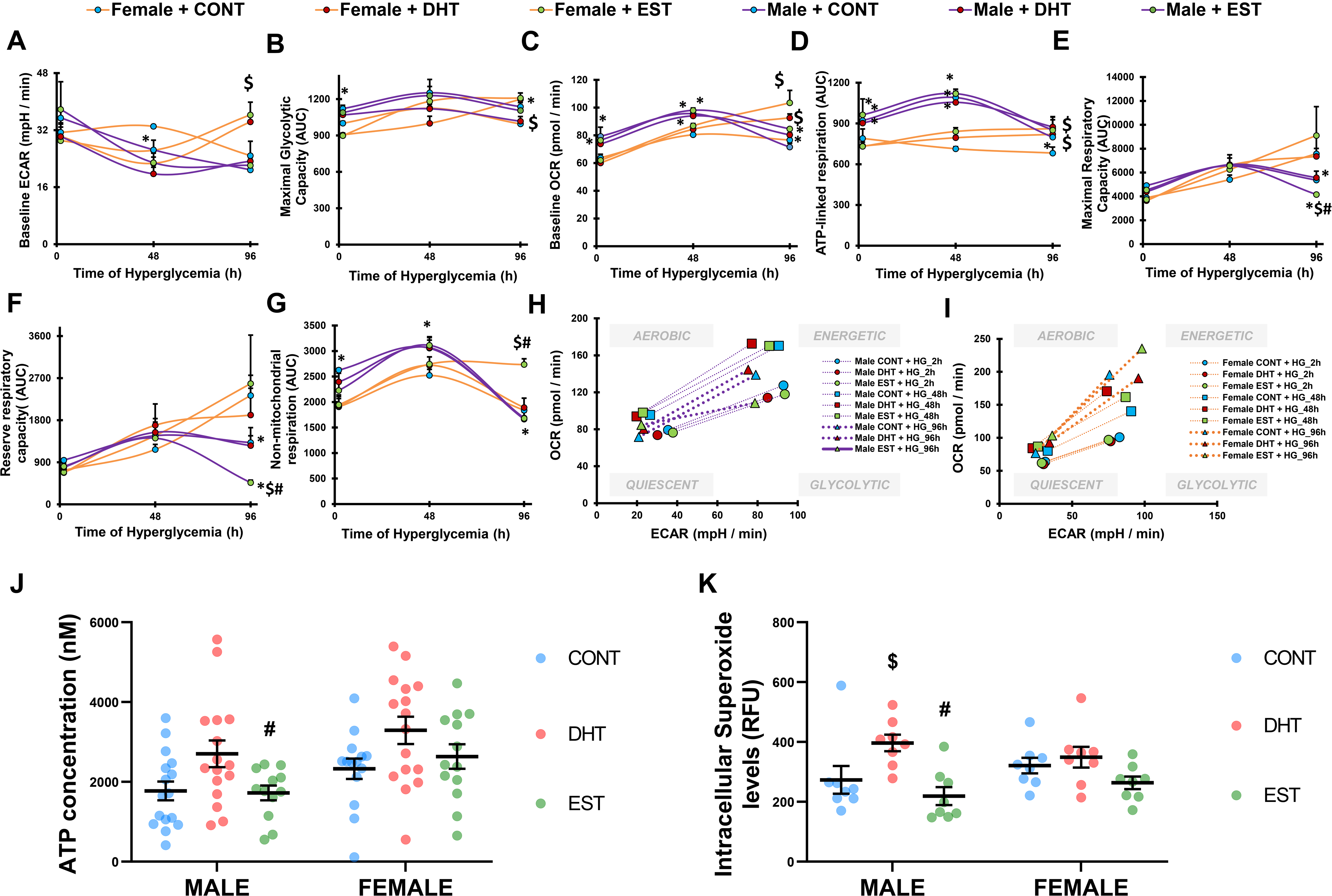
Effects of high glucose on the metabolism of male and female PTECs exposed to sex hormones. Male and female PTECs were serum starved and treated with vehicle (CONT), 100nM DHT or 100nM EST for 16h, prior to the exposure to high glucose. ECAR and OCR were monitored to assess glycolysis and mitochondrial function, respectively, at baseline and after 2h, 48h, and 96h of exposure to high glucose (25mM glucose; (n=2/sex; n=4-6/time point). DHT and EST concentrations were adjusted to 1nM during the high glucose treatment. The figure shows the evolution over time of key metabolic parameters: baseline glycolysis (A), maximal glycolytic capacity (B), basal respiration (C), ATP-linked respiration (D), maximal respiratory capacity (E), reserve respiratory capacity (F), and non-mitochondrial respiration (G). The evolution of the metabolic phenotype of male (H) and female PTECs (I) exposed to high glucose and sex hormones was visualized by plotting ECAR on the X axis and OCR on the Y axis. Intracellular levels of ATP (J) and superoxide ion (K) were also measured after 96h of exposure to high glucose. In panels A-G, a t test was used at each time point to assess statistical significance between the two groups. In panels J-K, group-to-group differences were determined using pairwise t tests for variables following a normal distribution, and Mann-Whitney tests for variables with a non-parametric distribution. *p<0.05 vs female PTECs; $p<0.05 vs CONT; #p<0.05 vs DHT. PTECs, proximal tubular epithelial cells; AUC, area under the curve; DHT, dihydrotestosterone; EST, estradiol; HG, high glucose; ECAR, extracellular acidification rate; OCR, oxygen consumption rate.

**Figure S5.**
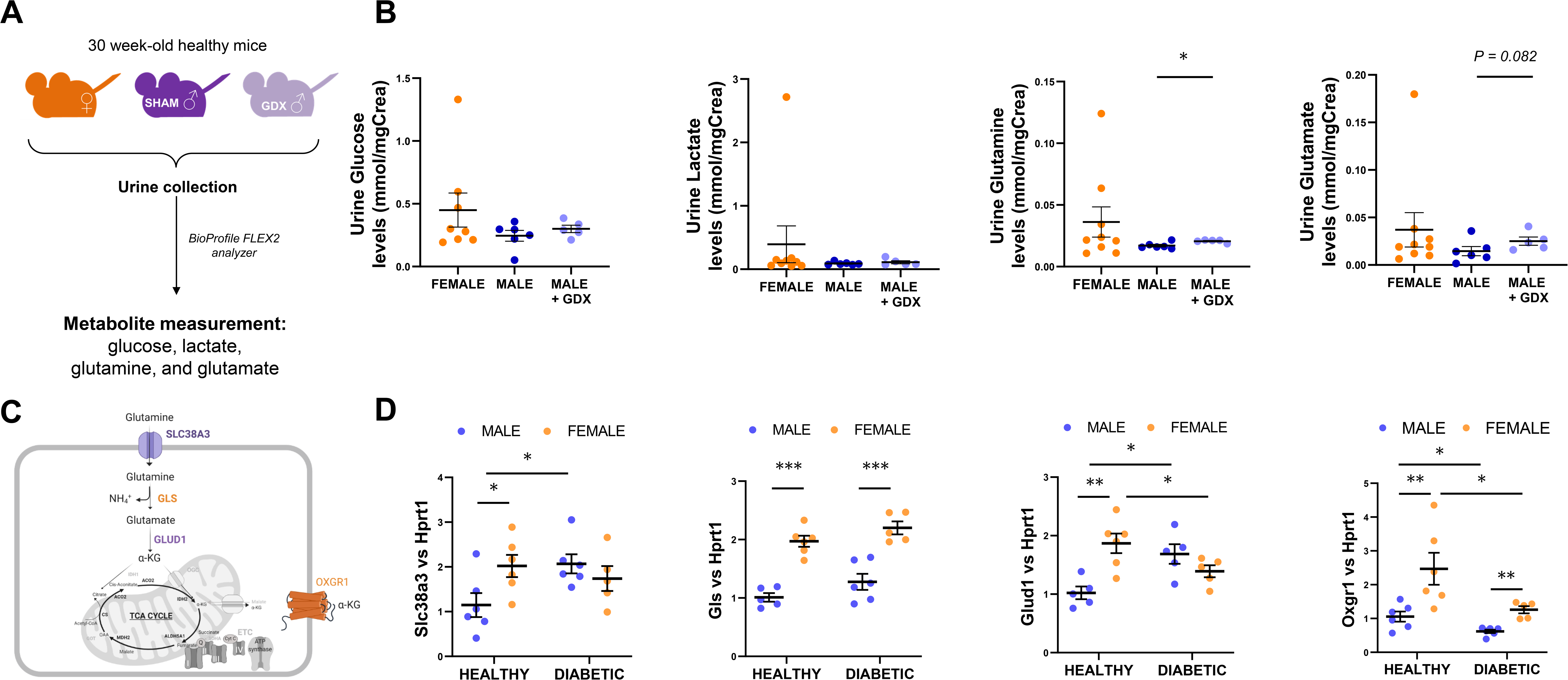
Sex-based differences on urine metabolites and kidney expression of genes related to glucose and glutamine metabolism *in vivo*. The urinary excretion of glucose, lactate, glutamine, and glutamate was measured in 30-week-old male mice previously subjected to gonadectomy (GDX). Metabolite levels in the urine of GDX male mice were compared to those in the urine of sham-operated males and intact females (A). Male mice displayed a reduction in the urinary levels of all four metabolites, in comparison to females, and this reduction was prevented by GDX (B) (n=5-8 animals/group). The kidney gene expression of *Slc38a3*, *Gls*, *Glud1*, and *Oxgr1* was measured in 16-week-old male and female, healthy and type 1 diabetic (Akita) mice, and normalized to Hprt1 (C,D) (n=5-6 animals/group). Group-to-group differences were determined using pairwise t tests for variables following a normal distribution, and Mann-Whitney tests for variables with a non-parametric distribution. *p<0.05; **p<0.01; ***p<0.001. TCA, tricarboxylic acid; α-KG, alpha-ketoglutarate; SLC38A3, sodium-coupled neutral amino acid transporter 3; GLS, glutaminase; GLUD1, glutamate dehydrogenase; OXGR1, alpha-ketoglutarate receptor 1; HPRT1, hypoxanthine-guanine phosphoribosyltransferase. The illustration in panel C was created with BioRender.com.

**Figure S6.**
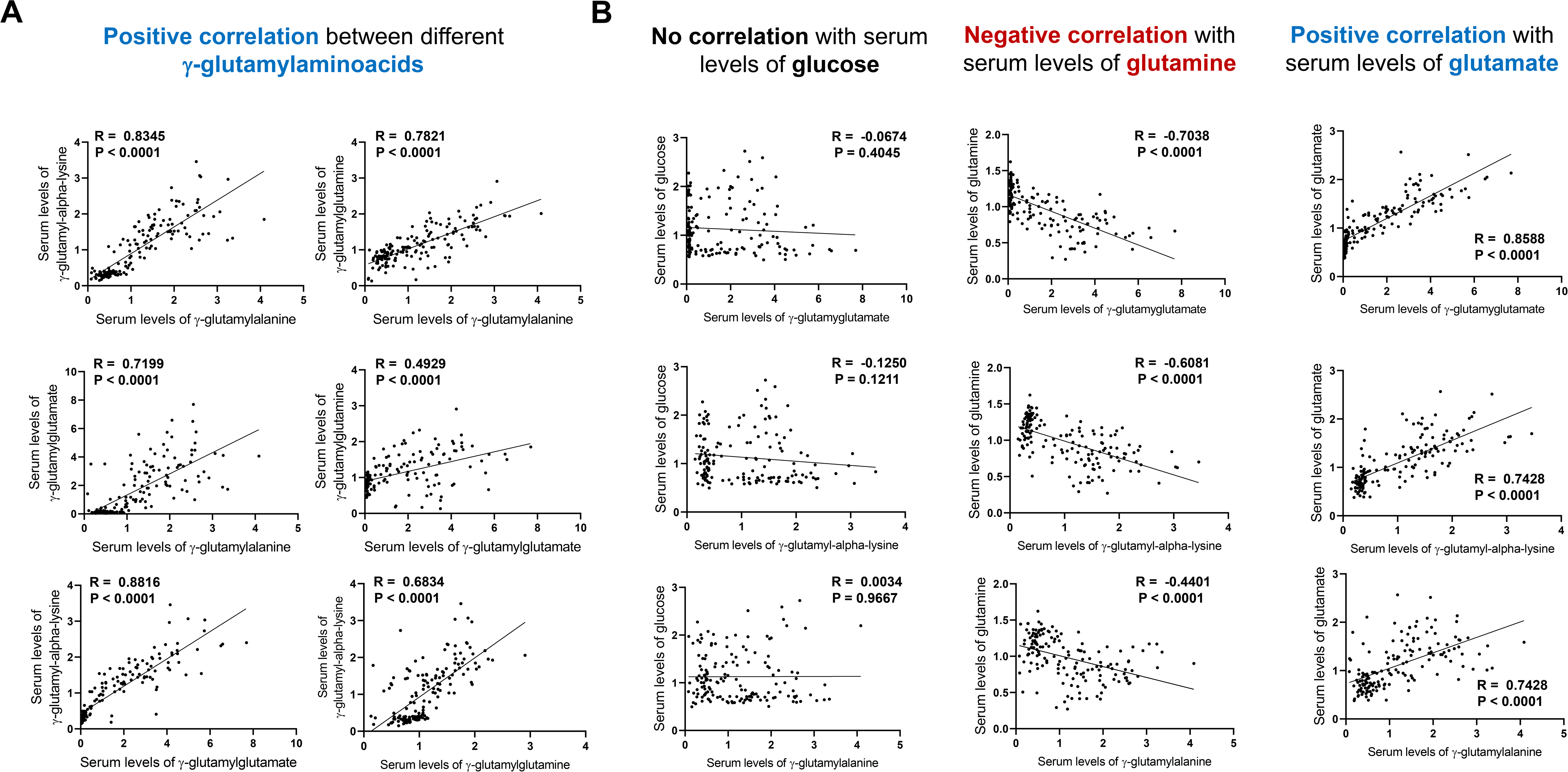
Representative correlations involving the circulating levels of γ-glutamyl-aminoacids, glucose, and glutamine in the human cohort. Pearson correlation was calculated for all possible pairs of circulating metabolites and correlation p-values were adjusted for multiple testing using the Benjamini-Hochberg method. The significance of each correlation was adjusted for an estimated 5% FDR. The scattered dot plots in panel A display the significant positive correlation between the circulating levels of 4 representative γ-glutamylaminoacids altered by sex and/or diabetes, namely γ-glutamyl-alpha-lysine, γ-glutamylalanine, γ-glutamylglutamine, and γ-glutamylglutamate. The serum levels of γ-glutamyl-alpha-lysine, γ-glutamylalanine, and γ-glutamylglutamate did not correlate with the levels of glucose. Instead, these metabolites showed a significant negative correlation with the levels of glutamine, and a significant positive correlation with the levels of unconjugated glutamate (B).

**Figure S7.**
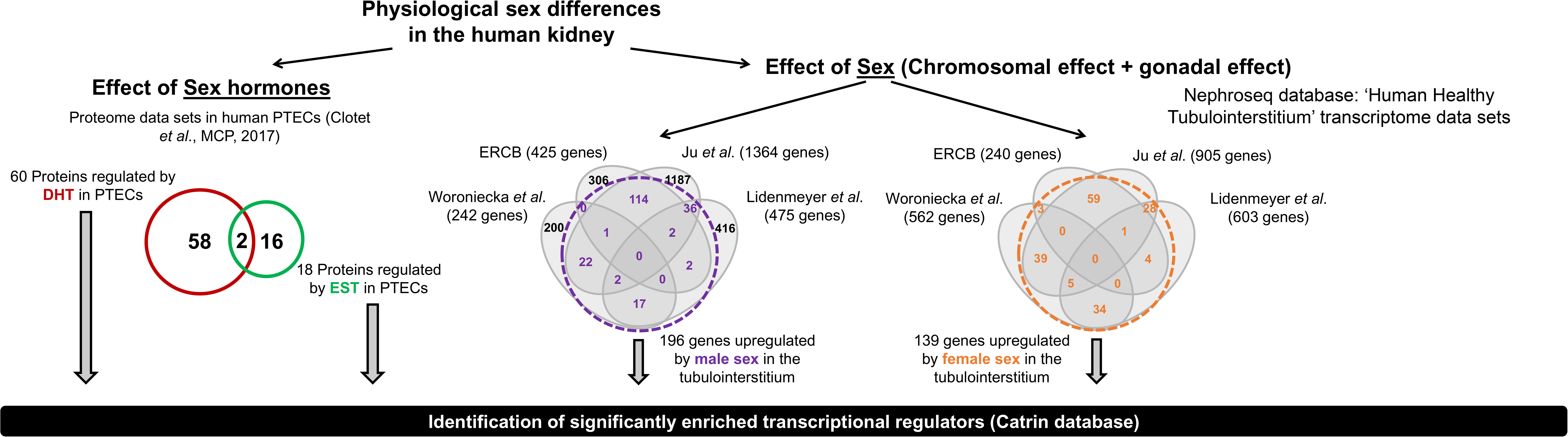
Identification of sex-biased genes and transcriptional regulators in the human kidney. To identify transcriptional mechanisms linked to the effect of sex hormones in the kidney tubule, we studied the 60 proteins upregulated by DHT, as well as the 18 upregulated by EST, in PTECs. To identify genes differentially expressed between male and female kidneys that could relate to our findings in tubular cells we analyzed publicly available gene expression data in Nephroseq database. Four data sets of the human healthy kidney tubulointerstitium including information about the sex of the sample were identified: Woroniecka *et al.*, Ju *et al*., Lindenmeyer *et al.,* and the European Renal cDNA Bank (ERCB, unpublished). The Venn diagrams depict the overlap between genes that showed a sex-biased expression with P<0.05 in each data set. Significantly altered genes showing the same direction of change in at least 2/4 data sets were retained. The analysis led to the identification of 196 genes significantly upregulated in the male tubulointerstitium and 139 genes significantly upregulated in the female tubulointerstitium.

**Figure S8.**
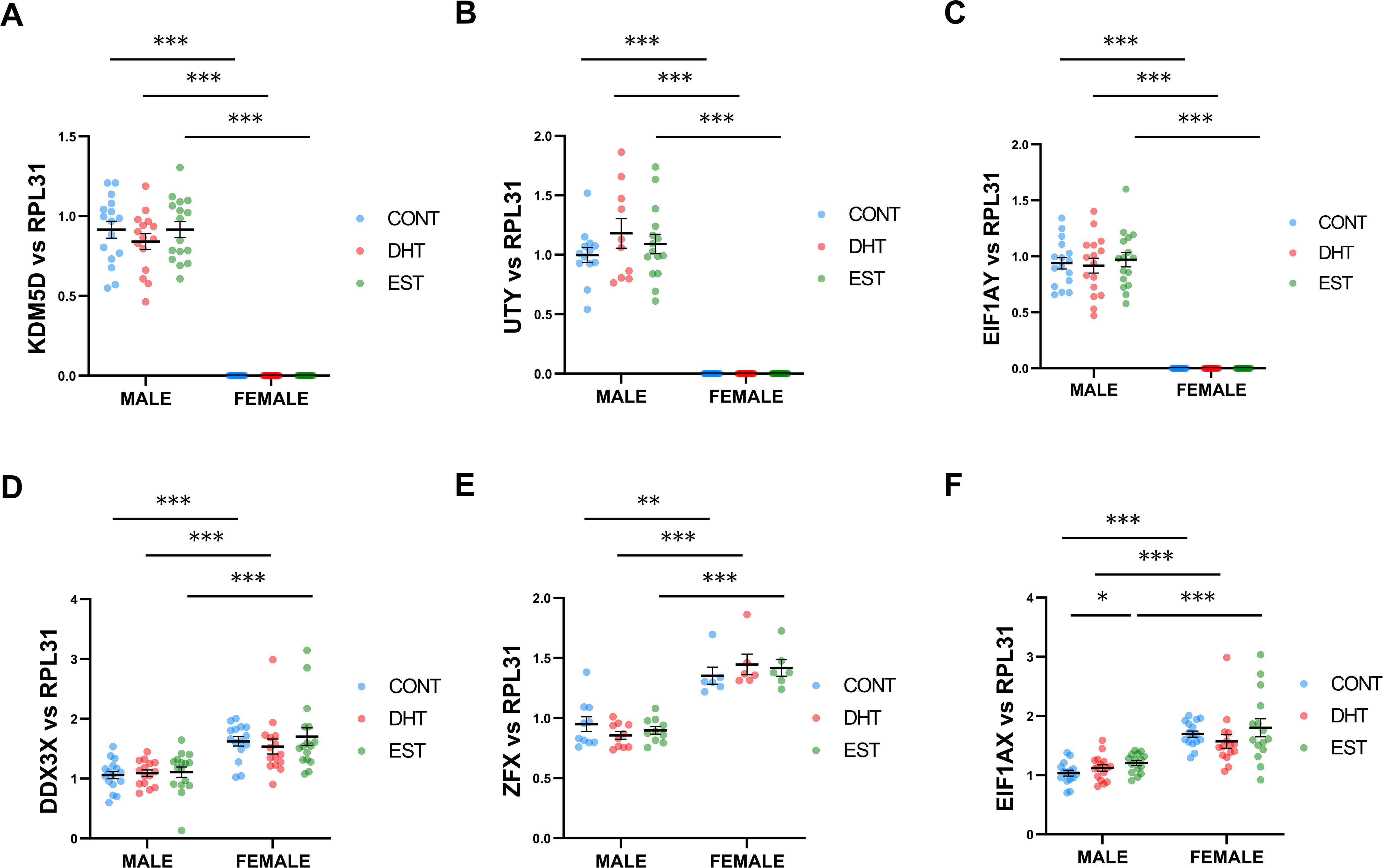
Sex-biased expression of Y-linked and X-escape genes in PTECs. Male and female PTECs were exposed to ethanol (CONT), 100nM DHT or 100nM EST for 16h (n=3/sex; n=4-6/treatment). The gene expression of KDM5D (A), UTY (B), EIF1AY (C), DDX3X (D), ZFX (E), and EIF1AX (F) was measured and normalized to RPL31. Group-to-group differences were determined using pairwise t tests for variables following a normal distribution, and Mann-Whitney tests for variables with a non-parametric distribution. *p<0.05; **p<0.01; ***p<0.001. PTECs, proximal tubular epithelial cells; DHT, dihydrotestosterone; EST, estradiol.

**Figure S9.**
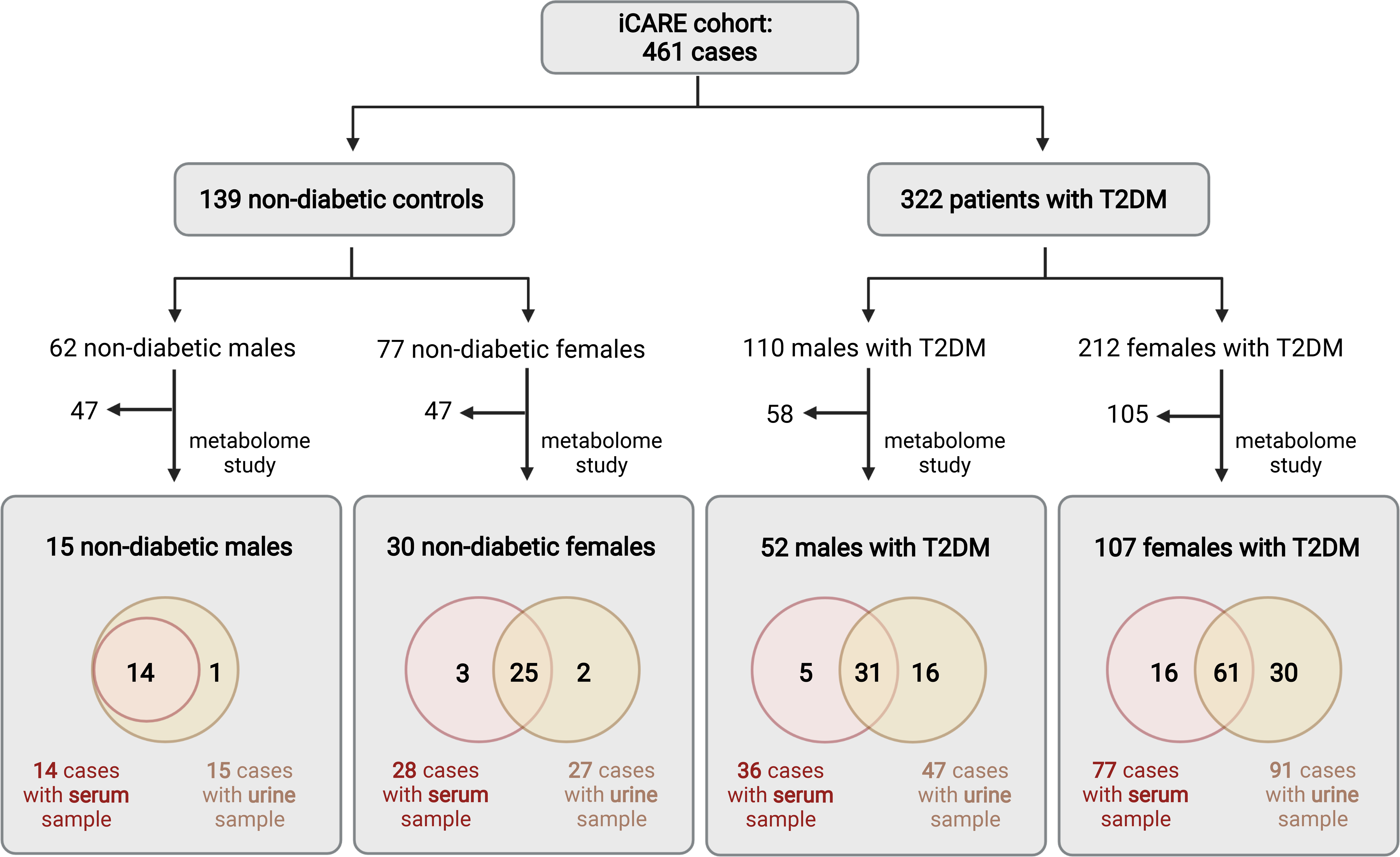
Serum and urine metabolome analysis in a subset of cases from the iCARE cohort. The flowchart illustrates the total number of cases encompassing the iCARE cohort, including non-diabetic males, non-diabetes females, males with T2DM, and females with T2DM. The subgroup of cases encompassing the metabolome study is also shown. T2DM, type 2 diabetes mellitus.

