## Supplemental Materials for "Cell Sex and Sex Hormones Modulate Kidney Glucose and Glutamine Metabolism in Health and Diabetes"

### SUPPLEMENTAL MATERIAL

#### SUPPLEMENTAL METHODS

##### Cell culture

*High glucose studies*

Male and female proximal tubular epithelial cells (PTECs) were grown and serum-starved as above. They were then treated with vehicle or sex hormones for 16h and exposed to Dulbecco's Modified Eagle Medium (DMEM) containing 25mM D-glucose, 4mM L-glutamine, and 1mM sodium pyruvate. Cells were studied at baseline and after 2h, 48h, and 96h of exposure to high glucose. To mimic decreased circulating sex hormone levels previously described in human DKD[1–5], and to prevent excessive cell death induced by high doses of sex hormones after a long exposure[6,7], DHT and EST concentrations were diminished to 1nM during the period of hyperglycemia. Co-treatment experiments were also conducted in male and female PTECs by simultaneously exposing them to sex hormones (100nM) and high (25mM) or normal glucose (5.55mM), for 16h or 24h.

*Pharmacologic inhibition studies*

Male and female PTECs were grown and serum-starved as described above, under normal glucose conditions. To study the effects of androgen receptor (AR) inhibition, male and female PTECs were pre-treated with flutamide (FLUT, F9397) or enzalutamide (ENZ, 1613 Axon MedChem) for 2h, prior to the exposure to CONT or DHT for 16h. To study the effects of KDM6A inhibition, male and female PTECs were treated with DMSO (vehicle), 2.5µM GSK-J4 (prototypical KDM6A inhibitor, ab144395 Abcam), or 2.5µM GSK-J5 (inactive control drug, ab144397 Abcam) for 48h. To study the effects of HNF4A inhibition, male and female PTECs exposed to CONT or DHT were co-treated with DMSO (vehicle) or 100nM BI6015 (prototypical HNF4A inhibitor, 375240 Sigma) for 16h.

**Induction of metabolic stress in kidney cells**

During the Seahorse assay, extracellular acidification rate (ECAR) and oxygen consumption rate (OCR) were recorded at baseline and after metabolic stress. To induce metabolic stress, 25µL of oligomycin, p-trifluoromethoxy carbonyl cyanide phenyl hydrazone (FCCP), 2-deoxyglucose (2-DG), and Rotenone + Antimycin A (Rot+AA) were sequentially injected into the microplate wells. After optimization, the following working concentrations were stablished for each drug: oligomycin: 1µM; FCCP: 0.3µM, 2-DG: 100mM; Rot: 1µM; AA: 1µM. Baseline glycolysis, maximal glycolytic capacity, and glycolytic reserve were assessed by calculating the area under the curve (AUC) from ECAR curves (**Fig.S1B**). Basal respiration, ATP-linked respiration, maximal respiratory capacity, reserve capacity, proton leak, and non-mitochondrial respiration were assessed by calculating the AUC from OCR curves (**Fig.S1C)**.

**Glucose uptake**

Glucose uptake in male and female PTECs was measured with a Glucose Uptake-Glo™ Assay Kit (Promega), following manufacturer instructions. Confluent cells were detached with 0.25% trypsin for 5min at 37ºC, and subsequently seeded in white 96-well microplates at a density of 15,000 cells/well in 100µL of DMEM complete media. After letting the cells adhere for 4-6h, PTECs were starved for 24h under normal glucose. Starvation DMEM media was removed, and cells were washed with PBS. Glucose uptake was initiated by incubating the cells with 50µL of 100µM 2-DG in PBS at room temperature (RT) for 10min. The reaction was neutralized by addition of the corresponding buffers (25µL of Stop Buffer and 25µL of Neutralization Buffer). The amount of 2-DG that entered the cells was then measured by exposing them to 100µL of 2DG6P

Detection reagent for 60min at RT, and subsequently recording the luminescent signal in a Cytation 5 plate reader (BioTek).

**Cell injury**

*Cellular oxidative stress and apoptosis*

Oxidative stress in male and female PTECs was assessed by measuring the intracellular levels of superoxide ion with the Cellular ROS Assay Kit (Red) (Abcam) following the manufacturer instructions. In turn, early apoptosis was assessed by measuring the levels of phosphatidylserine on the cell surface using the Apoptosis/Necrosis Assay Kit (Abcam). Confluent cells were cultured in the appropriate microplates and serum starved, as above. To measure superoxide ion levels, cells were stained with 100µL of ROS Red Working Solution for 45min at 37°C. Changes in fluorescence intensity were recorded at an Ex/Em=520/605nm in the plate reader. To measure the levels of phosphatidylserine at the cell surface, cells were stained with Apopxin Green Solution 1h at 37°C, and fluorescence was recorded at Ex/Em=490/525nm.

*Lactate dehydrogenase release*

Release of lactate dehydrogenase (LDH) in the cell supernatant was assessed as a marker of cell death in male and female PTECs using the CytoTox96® Cytotoxicity Assay (Promega) following the manufacturer instructions. Confluent cells were cultured in the appropriate microplates and serum starved, as above. To measure LDH release, 50µL of each cellular supernatant were loaded into different wells of a fresh 96-well flat clear bottom microplate. Fifty microliters of CytoTox96® Reagent were then added to each sample aliquot, and microplates were incubated for 30 min at RT and protected from the light. The reaction was terminated by adding 50µL of Stop Solution, and the absorbance signal (proportional to the number of lysed cells) was recorded at 490nM in the plate reader.

#### Cell viability

##### *Cellular ATP levels*

Intracellular levels of ATP in male and female PTECs were measured with a CellTiter-Glo 2.0 Assay Kit (Promega), following manufacturer instructions. To measure ATP levels, cells were washed with PBS and exposed to 100µL of CellTiter-Glo 2.0 Reagent (Promega). Contents were then mixed for 2min on an orbital shaker to induce cell lysis. Plates were then incubated at RT for 10min to stabilize the luminescent signal, which was recorded in the plate reader.

##### *MTT assay*

Cell viability was assessed in male and female PTECs using the MTT assay (Sigma), following manufacturer instructions. Serum-starved PTECs were exposed to the treatment of interest. After each treatment, 10µL of MTT labeling reagent were added into each well and cells were incubated at 37$^{o}$C and 5%$CO_2$ for 4h. After this period, 100µL of Solubilization buffer were added and cells were kept overnight in the same humidified atmosphere. Endogenous levels of formazan (generated through cleavage of MTT tetrazolium salt by enzymes of the endoplasmic reticulum of viable cells) were then measured at 550nM.

#### Cell metabolome

*Sample preparation*

Male and female PTECs were grown on 6-well plates and serum starved as described above. The steady-state intracellular metabolome was then determined after 16h of exposure to normal glucose starvation conditions. After collecting the cell supernatant, intracellular metabolites were extracted by adding 1mL of extraction solvent (80:20 mixture of methanol:water) into each well. With the plates on dry ice, the adherent material was triturated, collected into an Eppendorf tube, and stored at -80$^{o}$C. Cell lysate collection was followed by 3 cycles of freeze-thawing in dry ice (to shift sample temperature between -80$^{o}$C and -20$^{o}$C. The insoluble material from each collected sample was then precipitated at full speed for 5min. The resulting pellet was dried at RT and used for total RNA quantification using the Quant-iT Ribogreen assay (Invitrogen). In turn, the metabolite extract was dried under high purity nitrogen gas (turbovap) and resuspended with the appropriate volume of buffer (0.5µL of LC-MS grade water to 1µg of RNA) based on the total RNA levels measured in the corresponding pellet. Heavy-labeled (13C/15N) reference metabolites were spiked into each reconstituted sample as appropriate for quantitation. Heavy-labeled reference metabolites were acquired in the form of metabolite extract from yeast labeled to 99% by 13C-glucose and 15N-ammonia. A mock plate with no cells and the same volume of media was processed in parallel to the study plates, to obtain reference samples for determination of background metabolite signals.

*Liquid chromatography-mass spectrometry (LC-MS)*

Cellular metabolites were measured by injecting 2µL of sample in full scan MS1 mode using an Agilent 6550 qToF mass spectrometer coupled to an Agilent 1290 binary pump UPLC system. Most polar metabolite analytes presented here were measured using an Agilent ZORBAX Extend-C18 1.8 µm, 2.1 mm X 150 mm reverse phase chromatography using tributylamine as an ion paring agent as previously described[8]. The Agilent 6550 qToF was fitted with a dual AJS ESI source and an iFunnel with a gas temperature set to 150°C at 14L/min and 45psig. Sheath gas temperature was set to 325°C at 12L/min. Capillary and nozzle voltages were set to 2000V. Funnel conditions were changed from default to -30V DC, high pressure funnel drop -100V and RF voltage of 110V, low pressure funnel drop -50V and RF voltage of 60V.

A few small polar analytes (lactate, citrate, isocitrate, malate, fumarate, succinate, α-KG, 2-hydroxyglutarate, itaconic acid) were analyzed using a specialized method using an ACQUITY UPLC HSS T3 Column, 100Å, 1.8 µm, 2.1 mm X 150 mm column (Waters) kept at 45°C. This method used an isocratic gradient of 99.5% buffer A (10mM formic acid buffer pH 2.9 in water) and 0.5% Buffer B (Acetonitrile) held for 8 minutes. The column was then washed with 99% buffer B for 2 minutes and re-equilibrated with 0.5% buffer B for 2 minutes. Source conditions for this method were same as for ion-paired reverse phase above, however, default iFunnel conditions were used:  -50V DC, high pressure funnel -150V drop and RF voltage of 200V, low pressure funnel drop -100V and RF voltage of 100V.

Metabolite annotation in full scan data was achieved by matching exact mass and retention time to an in-house database. The retention time and exact mass database were prepared by analyzing a collection of neat standards using the chromatographic methods described above and confirming retention times by MS/MS fragmentation of neat standards.

*Metabolome data analysis*

Metabolome data was extracted directly from .d folders and integrated in profile mode using an R-based software package developed by the Rosebrock Lab; ChromXtractorPro (personal correspondence K. Laverty and A. Rosebrock,). The metabolites whose intensity in all the study samples fell at or below their intensity in the blank (consisting of resuspension buffer only) were excluded from further analyses. Next, the integrated light (L) intensity of each metabolite was normalized to the intensity of its internal heavy (H) standard. The L/H ratio minimized the potential stochastic variation in the signal produced by the instrument due to changes in humidity and/or temperature, enabling the relative quantitation and comparative analysis of each metabolite. The analysis enabled the detection of 158 intracellular metabolites (**Table S4B**). Significant group-to-group differences in metabolite levels were determined using unpaired two-tailed Student t tests, and the p-values were subsequently adjusted using the p.adjust() function in R for an estimated 5% false discovery rate. Among the metabolites quantified with confidence, 82 metabolites were significantly altered by cell sex (**Table S4C**).

**Animal studies**

*Surgical castration*

To evaluate the role of male and female sex hormones in the kidney levels of key transcriptional regulators, 10-week-old C57BL/6 healthy male and female mice were subjected to gonadectomy (GDX) and ovariectomy (OVX), respectively. After one week of rest, GDX or OVX was performed on male or female mice via a scrotal or abdomen midline incision, respectively. Mice were injected with buprenorphine and meloxicam subcutaneously before surgery and were deeply anesthetized with isoflurane during surgery. Sham control surgeries were also performed on male and female mice. Sham surgical procedures were the same as for GDX and OVX animals but without removing the gonads. Animals were followed for 10 weeks before being sacrificed.

To assess the effect of androgens on the urine excretion of glucose, lactate, glutamine, and glutamate, we studied the effect of GDX in a different colony of C57BL/6 10-week-old healthy male mice, as previously described[9]. Briefly, mice were anesthetized intraperitoneally with ketamine (75mg/kg) and medetomidine (1mg/kg), and after performing a vertical incision in the middle of the scrotal sac, the inner sacs, testes and epididymis were exposed. The epididymis was ligated with 4-0 stitches and testes were removed. Mice were recovered from anesthesia by subcutaneous atipamezol injection (1mg/kg). Intraperitoneal buprenorphine (0.5mg/kg) was administered as anti-inflammatory. To obtain the corresponding sham-operated groups, mice were anesthetized and the scrotal sac was exposed but not perforated. Testes were then returned to the original position in the perineal region and recovery from anesthesia was performed as in GDX animals. These mice were followed for 20 weeks and compared to sham-operated males and intact females of the same age.

**Targeted metabolite measurements**

Levels of glucose, lactate, glutamine, and glutamate were measured in cell supernatants, plasma and urine using a BioProfile® FLEX2™ Automated Cell Culture Analyzer (Nova Biomedical). At the end of each *in vitro* experiment, cell media from 6-well plates were collected and prepared for metabolite measurements by centrifugation at 2,000G for 10min at 4ºC, to remove cell debris. Treatment media from adjacent empty wells were subjected to the same processing and used as reference to calculate relative metabolite levels linked to cell metabolism. FBS, complete media (10% FBS) and starvation media (0% FBS) were employed as quality controls. Whereas plasma metabolite levels are expressed in mmols/L, urine metabolite levels were normalized to creatinine concentration and expressed in mmol/mg Crea. Urine creatinine was measured using a Colorimetric Creatinine Assay Kit (Abcam, ab204537) following manufacturer instructions.

**Gene expression**

RNA was extracted from PTEC cell pellets or from 30-50mg of mouse kidney cortex using the RNAeasy Mini Kit (Qiagen). After quantifying RNA concentration in a Nanodrop instrument (Thermo), 300-700ng of RNA were retrotranscribed to cDNA using the High-Capacity cDNA Reverse Transcription Kit (Applied Biosystems). For the *in vitro* experiments, male and female PTECs were treated with vehicle, 100nM DHT, or 100nM EST for 16-24h under normal glucose conditions. In these cells, gene levels of AR, KDM6A, ZFX, DDX3X, HNF4A, ZFY, EIF1AY, RPS4Y1, UTY, DDX3Y, and KDM5D were measured by real-time quantitative PCR using a Power SYBR® Green PCR Master Mix reagent (Applied Biosystems) and normalized to HPRT1 or RPL31. The fluorescent signal was measured in a StepOnePlus System (Applied Biosystems) for 96-well plates, and in a LightCycler® 480 Instrument II (Roche) for 384-well plates. For the *in vivo* experiments, gene levels of *Slc38a3*, *Gls*, *Glud1*, *Oxgr1, Kdm6a, Hnf4a, Zfy, Eif1ay, Uty, and Ddx3y* were measured and normalized to *Hprt1* or *Rpl31*. All primer sequences employed in this study are summarized in **Table S7**.

**Human serum and urine metabolomics**

*Sample preparation for metabolomics*

The 155 serum and 180 urine samples of the iCARE cohort were prepared using the automated MicroLab STAR® system (Hamilton Company). Recovery standards were included in the preparation process for quality control (QC) purposes. In each sample and standard, proteins were precipitated in absolute methanol for 2min through vigorous shaking (Glen Mills GenoGrinder 2000) and subsequent centrifugation. This step enabled the removal of protein, the dissociation of small molecules bound to proteins or trapped in the precipitated protein matrix, and the recovery of chemically diverse metabolites. The extract was then divided into 5 aliquots.

Aliquots 1 and 2 were analysed using two separate reverse phase (RP)/UPLC-MS/MS methods with positive ion mode electrospray ionization (ESI). Aliquot 3 was used for analysis by RP/UPLC-MS/MS with negative ion mode ESI. Aliquot 4 was analyzed by HILIC/UPLC-MS/MS with negative ion mode ESI. Aliquot 5 was kept at -80ºC as backup sample. In each aliquot, the organic solvent was removed by placing the sample briefly on a TurboVap® (Zymark). Extracts were stored under nitrogen until subsequent mass spectrometry analysis.

*Ultrahigh Performance Liquid Chromatography-Tandem Mass Spectroscopy (UPLC-MS/MS)*

The metabolome was measured in the serum and urine sample aliquots following an untargeted UPLC-MS/MS approach (Metabolon Inc, Durham, North Carolina). All four methods employed were performed using a Waters ACQUITY ultra-performance liquid chromatography (UPLC) and a Thermo Scientific Q-Exactive high-resolution mass spectrometer interfaced with a heated electrospray ionization (HESI-II) source and Orbitrap mass analyzer operated at 35,000 mass resolution. After drying, sample extracts were reconstituted in the appropriate solvents that corresponded to each of the four methods. Internal standards at fixed concentrations were previously added into each of the reconstitution solvents to ensure injection and chromatographic consistency.

Aliquot 1 was analyzed using acidic positive ion conditions, optimized via chromatography for more hydrophilic compounds. In this method, the extract was loaded in a C18 column (Waters UPLC BEH C18-2.1x100 mm, 1.7 µm) and eluted by gradient using water and methanol, containing 0.05% perfluoropentanoic acid (PFPA) and 0.1% formic acid (FA). Aliquot 2 was also analyzed using acidic positive ion conditions, but this method had been chromatographically optimized for more hydrophobic compounds. This extract was eluted by gradient from the same C18 column using water, methanol, acetonitrile, 0.05% PFPA and 0.01% FA and was operated at an overall higher organic content. Aliquot 3 was run on basic negative ion mode using a separate C18 column. The basic extracts were gradient eluted from the column using methanol and water, with 6.5mM ammonium bicarbonate at pH 8. Aliquot 4 was analyzed by negative ionization following elution from a HILIC column (Waters UPLC BEH Amide 2.1x150 mm, 1.7 µm) using a gradient consisting of water and acetonitrile with 10mM Ammonium Formate pH 10.8. The spectra analysis alternated between MS and data-dependent MSn scans based on dynamic exclusion. The scan range was slightly different between methods but covered a mass to charge ratio (m/z) of 70-1,000.

*Metabolite identification*

Raw data files from the MS runs were extracted, peak-identified, and processed for QC using Metabolon's hardware and software. Compounds were identified by comparison against the Metabolon library of >3,300 commercially available purified standards. Compounds were also compared with additional entries that were created for recurrent unknown entities in the mass spectra. This library contains authenticated standard molecules with their m/z ratio, retention time index (RI), and chromatographic data (including MS/MS spectra). Biochemical metabolite identifications were based on 1) accurate mass match to the library +/- 10ppm; 2) RI of the detected metabolite within a narrow RI window of the proposed identification; and 3) the MS/MS forward and reverse scores between the ions present in the experimental spectrum and the ions present in the library spectrum.

Curation and QC processes were conducted by Metabolon data analysts to ensure accurate and consistent identification of true chemical entities. Chemicals identified as system artifacts, miss-assignments, and background noise were excluded from further analyses. Proprietary visualization and interpretation software was used to confirm the consistency of peak identification among the various samples. For each sample, library matches for each compound were checked and corrected if necessary.

*Metabolite quantification and data normalization*

After curation, peak quantification for each metabolite was preformed using the area-under-the-curve. Since the samples were analyzed in multiple days, data normalization was performed to correct variation emerging from inter-day tuning differences on instrument performance. Specifically, each compound was corrected in run-day blocks by registering the medians to equal one (1.00) and normalizing each data point proportionately.

**Bioinformatics**

*Determination of sex-biased kidney genes in the human kidney tubulointerstitium*

To identify genes differentially expressed between male and female kidneys that could relate to our findings in tubular cells we analyzed publicly available gene expression data in Nephroseq database version 4 (URL: https://www.nephroseq.org/resource/main.html). Eleven studies were retained after setting the 'Sex' comparison (male vs female) as a primary filter. Among those, we identified 4 data sets of the human healthy kidney tubulointerstitium: Woroniecka *et al.*,[10] Ju *et al*.,[11] Lindenmeyer *et al.,*[12] and the European Renal cDNA Bank (ERCB, unpublished, URL: https://www.ekfs.de/en/scientific-funding/currently-funded-projects/european-renale-cdna-bank-kroner-fresenius-biopsiebank, data extracted from the latest update: 2018/04/01). An unbiased comparison of all overrepresented and underrepresented genes in each of these 4 data sets was performed. Genes differentially expressed between sexes with $P<0.05$ and showing the same direction of change in at least 2/4 data sets were retained. The analysis led to the identification of 196 genes significantly upregulated in the male tubulointerstitium and 139 genes significantly upregulated in the female tubulointerstitium (**Table S5, Fig.S8**). These two lists of genes were used to identify transcriptional regulators linked to male or female sex, as described below.

*Prediction of sex-specific transcriptional regulators of human kidney signatures*

To identify transcriptional mechanisms linked to the effect of sex hormones in the kidney tubule, we studied computationally which regulators are predicted to target the 60 proteins upregulated by DHT, as well as the 18 upregulated by EST, in PTECs[13]. To account for sex chromosome effects on the kidney tubule, we studied which regulators are predicted to target the 196 genes significantly upregulated in the male tubulointerstitium and the 139 genes significantly upregulated in the female tubulointerstitium (**Table S5, Fig.S8**). We queried these 4 lists of proteins and genes in CATRIN ver. 1.0.6.2, a transcriptional regulator database that integrates the findings of 15 stand-alone transcriptional regulator databases (http://ophid.utoronto.ca/Catrin). Regulators of the genes of interest were identified using CATRIN (excluding TF2DNA.experimental) and hypergeometric test was performed to identify which molecules significantly regulated the lists of genes of interest. P-value correction was performed using the Benjamini-Hochberg method. These analyses were run in R 4.0.3. The transcriptional regulators significantly enriched among sex-biased genes ($Q<0.05$), among proteins upregulated by DHT ($Q<0.05$) and/or among proteins regulated by EST ($P<0.05$), were further studied. The analysis of our sexually dimorphic kidney signatures led to the identification of 423 regulators linked to DHT, 129 regulators linked to EST, 608 regulators linked to male sex, and 818 regulators linked to female sex (**Fig.7-8, Table S6**).

#### Statistical analysis

*Experimental studies*

The statistical analysis of our *in vitro* and *in vivo* data was performed using GraphPad Prism v9.1.2. Normality of each study variable was assessed by using a Shapiro-Wilk test. Group-to-group differences were determined using pairwise t tests for variables following a normal distribution, and Mann-Whitney tests for variables with a non-parametric distribution. For all the analyses, $P<0.05$ was considered statistically significant. For cell metabolome data, significant group-to-group differences in metabolite levels were determined using two-tailed Student t-tests adjusted for an estimated 5% false discovery rate (FDR).

*Human studies*

Normalized serum and urine metabolome data was analyzed using R[14] version 4.0.2. Data from 65 metabolites of interest in serum and 19 metabolites in urine was analyzed using three statistical methods. The metabolites were selected based on their direct connection to glucose and glutamine metabolism. First, group-to-group differences in the levels of each metabolite were calculated using pairwise Mann-Whitney tests and p-values were adjusted for multiple testing using the Benjamini-Hochberg method. The levels of each metabolite were then represented as median (interquartile range) using box plots (Graphpad Prism v9.1.2). Second, Pearson correlation was calculated for all pairs of metabolites and correlation p-values were adjusted for multiple testing using the Benjamini-Hochberg method. Specifically, the significance of each correlation was adjusted for an estimated 5% FDR. Third, the interaction between the levels of each metabolite and our key factors of interest (sex, diabetes, or both) was evaluated through linear regression analyses using the lm function. The significance of the linear regression models was set at $P<0.05$.

#### SUPPLEMENTAL TABLES

**Table S1. Comparative analyses of serum metabolites in the human cohort**

**Table S2. Interaction analyses to evaluate the contribution of sex and diabetes to serum and urine metabolite levels**

**Table S3. Correlation analysis between serum metabolite levels**

**Table S4. Intracellular metabolites quantified and differentially altered by cell sex in PTECs**

**Table S5. Sex-biased genes in the kidney tubulointerstitium of healthy human subjects**

**Table S6. Transcriptional regulators enriched among genes and proteins with sex-biased renal expression**

**Table S7. Primer sequences**

#### SUPPLEMENTAL MATERIAL REFERENCES
