## Supplementary material for "Cell Sex and Sex Hormones Modulate Kidney Glucose and Glutamine Metabolism in Health and Diabetes": Table S7

**Table S7. Primer sequences used for Real Time qPCR analysis.**

| **Species** | **Gene** | **Gene name** | **Forward primer (5' → 3')** | **Reverse primer (5' → 3')** |
| --- | --- | --- | --- | --- |
| Human | Androgen receptor | **AR** | TACGGGGACATGCGTTTGGA | TTTCTGCTGGCGCACAGGTA |
| Human | ATP-dependent RNA helicase DDX3X | **DDX3X** | TTGCAGTGGAAAATGCGCTC | TATAGCGCCCTTTGCTGGCT |
| Human | ATP-dependent RNA helicase DDX3Y | **DDX3Y** | TTTGCTGAAGCTTGTCAAAGGG | ACTTGCTCCTCCACTCTGTTT |
| Human | Eukaryotic translation initiation factor 1A, Y-chromosomal | **EIF1AY** | TGGACGATTGGAAGCATTGTG | AGTCCCGTAGACCAACCAAT |
| Human | Hepatocyte nuclear factor 4-alpha | **HNF4A** | CCTACACCACCCTGGAATTTG | GTTGAGGTTGGTGCCTTCT |
| Human | Hypoxanthine-guanine phosphoribosyltransferase | **HPRT1** | GCTGAGGATTTGGAAAGGGTGT | GGCCTCCCATCTCCTTCATCA |
| Human | Lysine-specific demethylase 5D | **KDM5D** | GGCTCCCTGCTACGATCACA | CCTCATTGTCAAACGGGTGTGT |
| Human | Lysine-specific demethylase 6A | **KDM6A** | TTGGCCCAGGTGACTGTGAA | TGGGCCACCAAGAACCCATT |
| Human | 60S ribosomal protein L31 | **RPL31** | GCCGTTCTGCCATCAACGAA | TTGAGCCTGGTGTCAATGCG |
| Human | Histone demethylase UTY | **UTY** | TGGAGGACCTAATCCAAGTTTATGA | TGCAGAAATTTCCTGAAGAGCA |
| Human | Zinc finger X-chromosomal protein | **ZFX** | GCCCGCACGTCCGTC | CGGCAGTGACAGGCGG |
| Mouse | Glutaminase | ***Gls*** | ACAACGTCAGATGGTGTCATGC | GCTTGTGTCAACAAAACAATGTTGC |
| Mouse | Glutamate dehydrogenase 1, mitochondrial | ***Glud1*** | GCACTCTGGCTTGGCCTACA | GGCAGCTGTTCTCAGGTCCA |
| Mouse | Hepatocyte nuclear factor 4-alpha | ***Hnf4a*** | CTACACCACCCTGGAGTTTG | ATGAATTGAGGTTGGCACCTT |
| Mouse | Hypoxanthine-guanine phosphoribosyltransferase | ***Hprt1*** | TGTTGTTGGATATGCCCTTG | AATGACACAAACGTGATTCAAA |
| Mouse | Lysine-specific demethylase 6A | ***Kdm6a*** | GGGAATCTGGAAAATTTTGTGGTGC | TGAGGCGGATGGTAATGGAGG |
| Mouse | 2-oxoglutarate receptor 1 | ***Oxgr1*** | GGGGAAGTCCCGTTTCTTTAAGC | GTGTCCTCTGTTCCCCGAACT |
| Mouse | 60S ribosomal protein L31 | ***Rpl31*** | CCCACTACCTGCTTAGAGCC | TTCTTTGAGTGCCCGAGGAG |
| Mouse | Sodium-coupled neutral amino acid transporter 3 | ***Slc38a3*** | TCCTTGTACCCCGAGACCCG | CCCCAGAACACTGCTCTCACA |
| Mouse | Histone demethylase UTY | ***Uty*** | TCACAGTCGAAGAGAAGAAGGC | TCTGATTCCACTTTTCCTTCAGC |
